# Sex differences in offspring risk and resilience following 11β-hydroxylase antagonism in a rodent model of maternal immune activation

**DOI:** 10.1101/2023.09.21.558903

**Authors:** Julia Martz, Micah A. Shelton, Laurel Geist, Marianne L. Seney, Amanda C. Kentner

**Author notes:** Corresponding author: Amanda Kentner Office.

## Abstract

Maternal immune activation (MIA) puts offspring at greater risk for neurodevelopmental disorders associated with impaired social behavior. While it is known that immune signaling through maternal, placental, and fetal compartments contributes to these phenotypical changes, it is unknown to what extent the stress response to illness is involved and how it can be harnessed for potential interventions. To this end, on gestational day 15, pregnant rat dams were administered the bacterial mimetic lipopolysaccharide (LPS; to induce MIA) alongside metyrapone, a clinically available 11β-hydroxylase inhibitor used to treat hypercortisolism in pregnant and neonatal populations. Maternal, placental, and fetal CNS levels of corticosterone and placental 11βHSD enzymes type 1 and 2 were measured 3-hrs post treatment. Offspring social behaviors were evaluated across critical phases of development. MIA was associated with increased maternal, placental, and fetal CNS corticosterone concentrations that were diminished with metyrapone exposure. Metyrapone protected against reductions in placental 11βHSD2 in males only, suggesting that less corticosterone was inactivated in female placentas. Behaviorally, metyrapone-exposure attenuated MIA-induced social disruptions in juvenile, adolescent, and adult males, while females were unaffected or performed worse. Metyrapone-exposure reversed MIA-induced transcriptional changes in monoamine-, glutamate-, and GABA-related genes in the ventral hippocampus of adult males, but not females. Taken together, these findings illustrate that MIA-induced HPA responses act alongside the immune system to produce behavioral deficits. As a clinically available drug, the sex-specific benefits and constraints of metyrapone should be investigated further as a potential means of reducing neurodevelopmental risks due to gestational MIA.

## Introduction

Exposure to inflammation during pregnancy can increase offspring susceptibility to neuropsychiatric disorders and dysregulated stress responses (**Han et al., 2021; Quagliato et al., 2021)**. Associated disorders, such as autism and schizophrenia, are characterized by a heterogeneous set of symptoms that include social and cognitive impairments which emerge in later life (**Estes & McAllister, 2016**). Moreover, these observed disruptions can be mirrored in laboratory animals by administering bacterial and viral mimetics during pregnancy (Bao et al., 2022). The detrimental effects of maternal immune activation (MIA) are mediated by a combination of immune and endocrine responses that affect the placenta and fetal brain **(Estes & McAllister, 2016; Quagliato et al., 2021)**. However, the precise mechanisms underlying these effects remain unclear.

MIA triggers the release of pro-inflammatory cytokines, such as interleukin (IL)-6 and IL-17A, along with other immunological factors that can influence fetal brain development by crossing the placenta and fetal blood brain barriers **(Wu et al., 2017; Choi et al., 2016; Hsiao et al., 2012; Stelzer et al., 2021; Iqbal et al., 2012**). However, like other forms of prenatal stress, MIA also activates the maternal hypothalamic-pituitary-adrenal (HPA) axis, potentially influencing offspring neural development and behavior through excess fetal exposure to glucocorticoids (**Glover et al., 2018; Wyrwoll & Holmes, 2012; Núñez Estevez et al., 2020**). During pregnancy, the maternal HPA axis is downregulated and 11β-hydroxysteroid dehydrogenase (11βHSD) enzyme types 1 and 2 act on the placenta to modulate glucocorticoid activity (**Neumann et al., 1998; Wyrwoll et al., 2011**). Specifically, 11βHSD2 inactivates corticosterone (cortisol in humans) by conversion to inactive 11-DHC (cortisone in humans). In contrast, 11βHSD1 converts inactive glucocorticoids to their active form (**Wyrwoll et al., 2011**). Inhibition of 11βHSD2 during pregnancy leads to dysregulated stress responses in offspring, suggesting a protective role of 11βHSD2 against excessive glucocorticoid exposure during fetal development (**Welberg et al., 2000**).

It has been proposed that chronic maternal restraint stress decreases 11βHSD2 activity, allowing elevated levels of glucocorticoids to pass through the placenta, enter the fetus, and disrupt brain development by affecting fetal glucocorticoid receptors (**Welberg et al., 2000 Mairesse et al., 2007; Jensen Peña et al., 2012**). While studies on this effect in maternal psychological stress models remain inconclusive (**Sze et al., 2022**), lipopolysaccharide (LPS)-induced MIA downregulates 11βHSD2 mRNA (**Straley et al., 2014; Núñez Estevez et al., 2020**), leading to disruptions in glucocorticoid metabolism and subsequent changes in glucocorticoid and corticotropin-releasing hormone (CRH) receptor expression in offspring (**Núñez Estevez et al., 2020; Connors et al., 2014; Zhao et al., 2020**). These central changes were accompanied by sex-specific alterations in social behaviors, including reduced social preference and impaired social discrimination. Although it is known that MIA activates the HPA axis, it is unknown to what extent this stress response contributes to the negative impact of MIA on fetal development (**Zhao et al., 2021**).

To investigate the influence of MIA-induced glucocorticoid synthesis on the placenta and fetal brain, the current study employed the clinically available drug metyrapone, a reversible inhibitor of cytochrome P450 11B1, mitochondrial (11β-hydroxylase), the enzyme that catalyzes the final step of cortisol synthesis in the adrenal cortex. This inhibitor is safely used in pregnant and lactating people, and in pediatric populations (**Azzola et al., 2020; Kersten et al., 2020; Bass et al., 2021; Duke et al., 2019; Lourenço et al., 2015; de Mingo et al., 2020**). Moreover, when used in pregnant laboratory rats, 11β-hydroxylase inhibitors have not impacted placental or offspring viability (live births and survival), sex ratio distribution of the litter, nor resulted in other signs of toxicity (**Leret et al., 2009; Ménard et al., 2014; Minck et al., 2019**). We hypothesized that attenuating the LPS-induced rise in plasma corticosterone with this 11β-hydroxylase inhibitor would prevent the placental, fetal, and postnatal disruptions that result following MIA challenge. To this end, we investigated the effect of LPS and MET on placental and fetal tissues, offspring behavior across development, and on brain transcriptional profiles.

## Materials and Methods

### Animals and Experimental Overview

Please see **Supplementary Methods** for more detailed descriptions of the study protocols and statistical analyses. Animal procedures were approved by the Massachusetts College of Pharmacy and Health Sciences (MCPHS) Institutional Animal Care and Use Committee and carried out in line with the Association for Assessment and Accreditation of Laboratory Animal Care. Experimental procedures and groups are presented in **Figure 1A** and **Supplementary Results Table 1. Supplementary Methods Table 1** provides additional methodological details **(Kentner et al., 2019)**.

**Figure 1.**
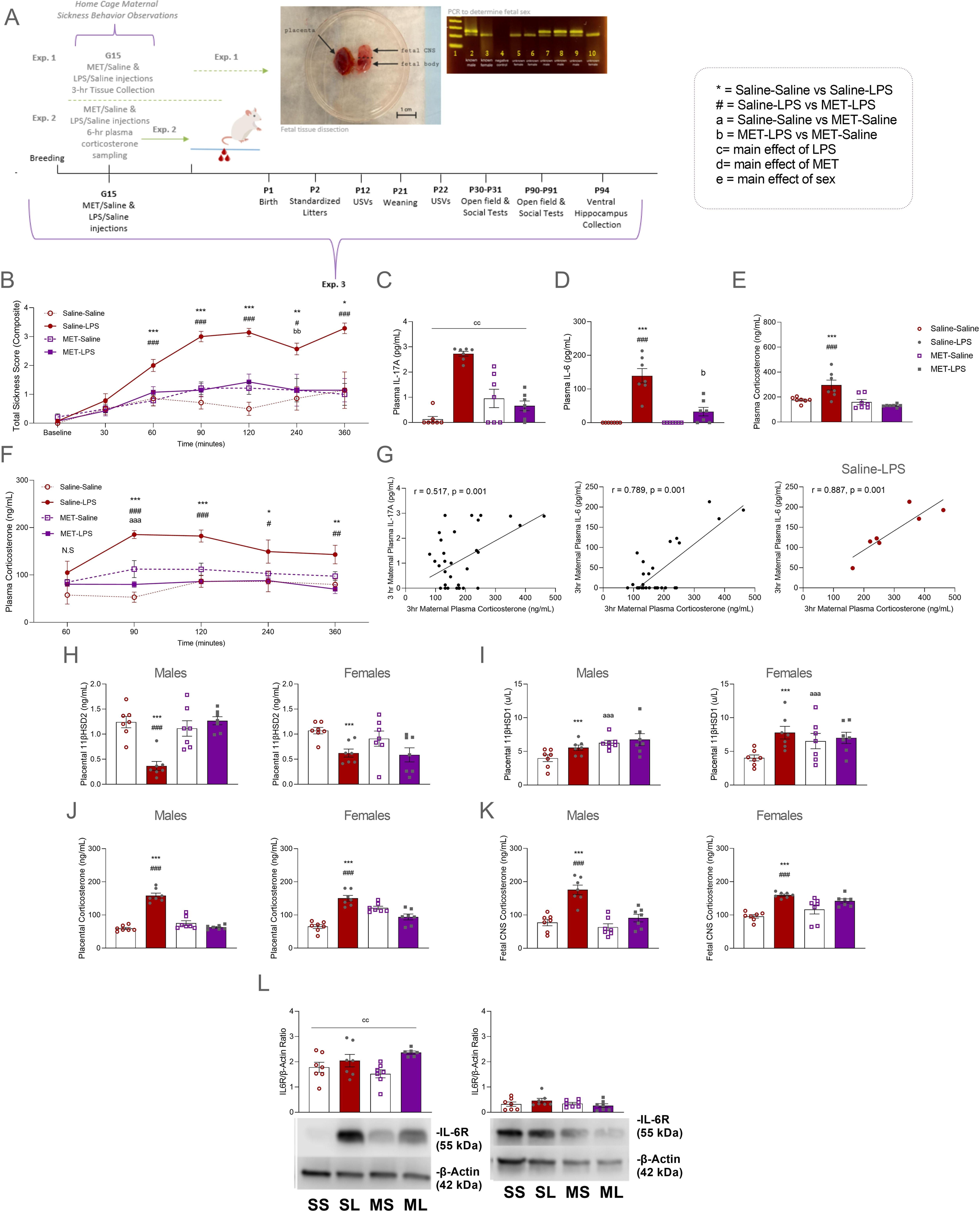
Effect of metyrapone (MET) on maternal immune activation (MIA)-induced sickness behaviors and 3-hr post LPS physiological response measures in dams, placenta, and fetal brain. A Timeline of experimental procedures. B Maternal sickness scores evaluated across a 6-hr period. Maternal sickness scores were elevated in Saline-LPS dams, and attenuated in MET-LPS dams, post MIA challenge. C Maternal plasma IL-17A concentrations were elevated by MIA. D Plasma IL-6 was elevated in MIA dams, which was modestly attenuated by MET. E MET protected against MIA induced elevations in maternal plasma corticosterone. F MET attenuated plasma corticosterone concentrations over a 6-hour period post G15 LPS challenge. G Maternal plasma corticosterone was positively correlated with maternal plasma IL-17A and maternal plasma IL-6 3-hours post MIA challenge. H Placental enzyme 11βHSD2 concentrations were reduced in Saline-LPS male and female placentas compared to the Saline-Saline groups 3-hours post MIA challenge. MET protected against decreased levels of 11βHSD2 in males only. I Placental enzyme 11βHSD1 was increased in Saline-LPS males and females compared to the Saline-Saline groups 3-hours post MIA challenge. Corticosterone was elevated in Saline-LPS male and female J placentas and K fetal brains compared to Saline-Saline groups, but not in MET-LPS compared to Saline-LPS placentas 3-hours post challenge. L MIA was associated with increased IL-6R in male fetal brains, irrespective of MET treatment. Data are expressed as mean ± SEM. ***p < 0.001, **p < 0.01, *p <0.05., Saline-Saline vs Saline-LPS; ^###^p < 0.001, ^##^p < 0.01, ^#^p <0.05, Saline-LPS vs MET-LPS; ^aaa^p < 0.001, ^aa^p < 0.01, ^a^p <0.05, Saline-Saline vs MET-Saline; ^bbb^p < 0.001, ^bb^p < 0.01, ^b^p <0.05, MET-LPS vs MET-Saline; ^cc^p < 0.01, main effect of LPS/Saline. LPS – lipopolysaccharide.

On the morning of gestational day (G) 15, Sprague Dawley rat dams were administered pyrogen-free saline (vehicle) or the 11β-hydroxylase inhibitor metyrapone (MET, 100 mg/kg, i.p). One hour later, dams were treated with pyrogen-free saline (vehicle), or 200 μg/kg (i.p.) of the inflammatory endotoxin lipopolysaccharide (LPS; Escherichia coli, serotype 026:B6; L-3755) to induce MIA.

#### Experiment 1

To validate MIA induction by LPS, sickness behaviors were evaluated as previously described (**Connors et al., 2014; Hayley et al., 2002;** N = 56). A subset of dams (n = 7) was euthanized by ketamine/xylazine (150 mg/kg/50 mg/kg, i.p.) 3 hours following the G15 LPS challenge. Maternal cardiac plasma samples were collected and stored for further validation of MIA by ELISA. Dams were perfused intracardially with a phosphate buffer solution (PBS) and the uterine horn removed. Placentas, whole fetal brains, and bodies were saved for later processing (**Bronson & Bale, 2014**). Genomic DNA was extracted from embryonic tissue using DNeasy Blood & Tissue Kits (QIAGEN, Valencia, CA, USA), according to manufacturer’s instructions. Sex was determined by the presence of a Sry band (protocol adapted from **Miyajima et al., 2009; Núñez Estevez et al., 2020; Figure 1A**). ELISAs were used to measure corticosterone levels of maternal plasma, fetal brains, and placenta (n = 7; Enzo Life Sciences, Farmingdale, NY), maternal plasma cytokines (IL-6 and IL-17A, Abcam, Waltham, MA), and placental 11 β-hydroxysteroid dehydrogenase (11βHSD1 and 11βHSD2, MyBioSource, San Diego, CA). Fetal brains were evaluated by Western blot (corticotropin releasing factor receptor 1 [CRFR1]; glucocorticoid receptor [GR], interleukin [IL]-6 receptor [IL-6R]; IL-17A and β-actin, Thermo Fisher Scientific, Waltham, MA).

#### Experiment 2

A second set of dams was used to validate that the 11β-hydroxylase inhibitor MET attenuated the sustained release of maternal plasma corticosterone in response to LPS (n = 5-7) as measured by ELISA.

#### Experiment 3

Following G15 drug treatments, a third group of dams (n = 10-11) continued pregnancies through until birth (postnatal day (P)1). A P4 maternal retrieval test took place (**Spencer et al., 2006)** and offspring were evaluated on several behavioral metrics, including isolated and social ultrasonic vocalizations (USVs; **from Bodi et al., 2016; Kentner et al., 2018; Van Segbroeck et al., 2017**), open field, social preference, and social discrimination tests, as previously described (**see Yan & Kentner, 2017; Connors et al., 2014; Zhao et al., 2021a; Núñez Estevez et al., 2020; Scarborough et al., 2020**). Adult ventral hippocampus was collected and stored at −80°C until western blot processing for GR and CRFR1 (n = 6-7) or RNA sequencing (n = 3). RNA was extracted using RNeasy Plus Micro kits (Qiagen, Valencia, CA, USA), RNA libraries were prepared and sequenced at Azenta Life Sciences (South Plainfield, NJ, USA). Illumina NovaSeq 6000 (Illumina, San Diego, CA, USA) was used to sequence the samples according to the manufacturer’s instructions using a 2×150bp Paired End (PE) configuration.

Friedman’s non-parametric version of the one-way repeated measures ANOVA was used to evaluate maternal sickness behavior. Kruskal-Wallis was used for follow-up tests. Fetal tissues were adequately powered to detect sex-differences, so three-way ANOVAs (Sex x G15 MET/Saline x G15 LPS/Saline) were utilized with litter as a covariate. Two-way ANOVAs were used as appropriate for all other measures (G15 MET/Saline x G15 LPS/Saline) unless the data were skewed (Shapiro-Wilks) then non-parametric Kruskal-Wallis tests were employed (see **Green & Salkind, 2005**). Because behavioral and adult tissue datasets were not powered to evaluate sex-differences directly, combined with the fact that we were interested in sex-specific effects, male and female animals were evaluated separately (**Ordoñes Sanchez et al., 2021**). Partial eta-squared (*n* ^2^) is reported as an index of effect size for ANOVAs (**Miles & Shevlin, 2001**). Pearson correlations were utilized to determine associations for prenatal and postnatal measures. All data are expressed as mean ± SEM. For RNA-sequencing, CLC Genomics Workbench v.23.0.2 software (QIAGEN Digital Insights, Aarhus, Denmark) was used for differential expression (DE) analysis. Our approach was to identify genes in which the effect of LPS was reversed by MET: 1) We first probed for effects of LPS by comparing expression of Saline-LPS animals to the Saline-Saline group; 2) We next compared expression in MET-LPS animals to the Saline-LPS group. Putting these differential expression analyses together, we could determine which genes were altered by LPS for which the effect was reversed by MET. Volcano plots for each brain region were constructed using the log2 fold and -log10 p-value change of expression in GraphPad Prism v.9.5.1. Heatmaps representing our expression data were created using the free browser program Morpheus (https://software.broadinstitute.org/morpheus). Genes with p < 0.05 and fold change > 1.3 were considered DE. RRHO2 was used as a threshold-free approach to determine the extent of concordance/discordance between transcriptomic datasets (**Cahill et al., 2018**).

## Results

### MET protected against MIA-induced sickness behaviors and blocked elevations in plasma corticosterone in dams

Consistent with previous studies, there was a significant time effect of LPS treatment on total sickness scores (Saline-Saline vs Saline-LPS; Friedman’s non-parametric test; *X*^2^(5) = 11.970, p = 0.001). Saline-LPS dams demonstrated robust sickness behaviors starting 60 minutes post LPS challenge, which was significantly attenuated by MET (Saline-LPS vs MET-LPS; p < 0.05; **Figure 1B**). Based on the skewness of the IL-17A data, a non-parametric analysis showed elevated cytokine levels following MIA (main effect LPS/Saline: (*X*^2^(1) = 8.669, p = 0.003; **Figure 1C**). Surprisingly, MET partially protected against MIA-induced elevations in plasma IL-6 (G15 MET/Saline by G15 LPS/Saline interaction: F(1, 24) = 18.437, p = 0.001, n_p_ ^2^ = 0.434; Saline-Saline vs Saline-LPS: t(12) = -6.518, p = 0.001; Saline-LPS vs MET-LPS: t(12) = 4.294, p =0.001; **Figure 1D**); this was only modest as they still had significantly higher concentrations of this cytokine than MET-Saline animals (p <0.05; **Figure 1D**). MET-Saline dams did not differ from Saline-Saline on this measure (p>0.05). Importantly, MET completely protected from the increased corticosterone concentrations induced by LPS at the 3-hr mark (G15 MET/Saline by G15 LPS/Saline interaction: F(1, 24) = 0.401, p = 0.004, n_p_ ^2^ = 0.302; Saline-Saline vs Saline-LPS: t(12) = - 2.942, p = 0.012; Saline-LPS vs MET-LPS: t(12) = 4.132, p = 0.001; **Figure 1E**). A repeated measures ANOVA confirmed MET had a sustained influence on the attenuation of MIA-induced plasma corticosterone when sampled across a six-hour period (time x G15 MET/Saline x G15 LPS/Saline interaction: F(7.38, 14) = 5.05, p = 0.001, n_p_ ^2^ = 0.444; Experiment 2; **Figure 1F**). **Supplemental Results Statistical Table 2** outlines correlational results for several prenatal measures with a selection of data displayed in **Figure 1G**. Together, these results suggest that blocking corticosterone synthesis with MET partially reverses the effects of MIA on maternal immune responses.

### MET affected placenta and fetal CNS tissues in a sex and MIA dependent manner

Three-hours post MIA, 11βHSD2 concentrations were significantly reduced by LPS in both sexes (Saline-Saline vs. Saline-LPS; p = 0.001; **Figure 1H**). Male animals pretreated with MET were fully protected against decreased 11βHSD2 levels (Saline-LPS vs MET-LPS: p = 0.001; **Figure 1H**). In contrast, placental 11βHSD1 was elevated in male and female Saline-LPS and MET-Saline groups compared to Saline-Saline (p = 0.001; **Figure 1I**).

Corticosterone concentrations were elevated by LPS in placental and fetal brain 3-hours following MIA (Saline-Saline vs. Saline-LPS; p = 0.001; **Figure 1J,K**). MET attenuated these effects (MET-LPS versus Saline-LPS: p = 0.001; **Figure 1J,K**). Interestingly, IL-6R was elevated in male fetal brains exposed to LPS (p = 0.01), irrespective of MET treatment, however there were no receptor level differences across groups in females (p > 0.05; **Figure 1L**). There were no effects of gestational LPS on CRFR1, GR, or IL-17A in male or female fetal brains 3-hrs post challenge (p > 0.05; **Supplemental Results Figure 1**).

### MET blocked MIA-induced behavioral changes in a sex-dependent manner

There were no group differences in the maternal retrieval test on P4. While there was a modest main effect of MET on the latency to retrieve the entire litter (p = 0.042), the post hoc test was non-significant (p > 0.05, **Supplemental Results Statistical Table 2**).

In the P12 neonatal isolation USV test, male and female MIA animals had a reduced number of syllables emitted during both the baseline and potentiation period (main effect of LPS: p = 0.05; **Figure 2A**). During the potentiation period, MET exposed males had a higher number of syllables/calls (p < 0.05; **Figure 2A**). Male and female MIA offspring also had an attenuated number of calls during the P22 social play test (Saline-Saline vs Saline-LPS: p = 0.001; **Figure 2B**); this effect was prevented in LPS males pre-treated with MET (Saline-LPS vs MET-LPS: p = 0.01; **Figure 2B**). There were no differences across groups for the mean duration of syllables for either the P12 USVs or the P22 social play induced-USVs (**Supplemental Results Figure 2A,B**). **Figure 2C** shows the top vocalization types for P12 and P22 USV tests. The total number of syllables emitted during the USV tests were negatively correlated to the severity of maternal sickness behaviors (**Figure 2D; Supplemental Results Statistical Table 2**).

**Figure 2.**
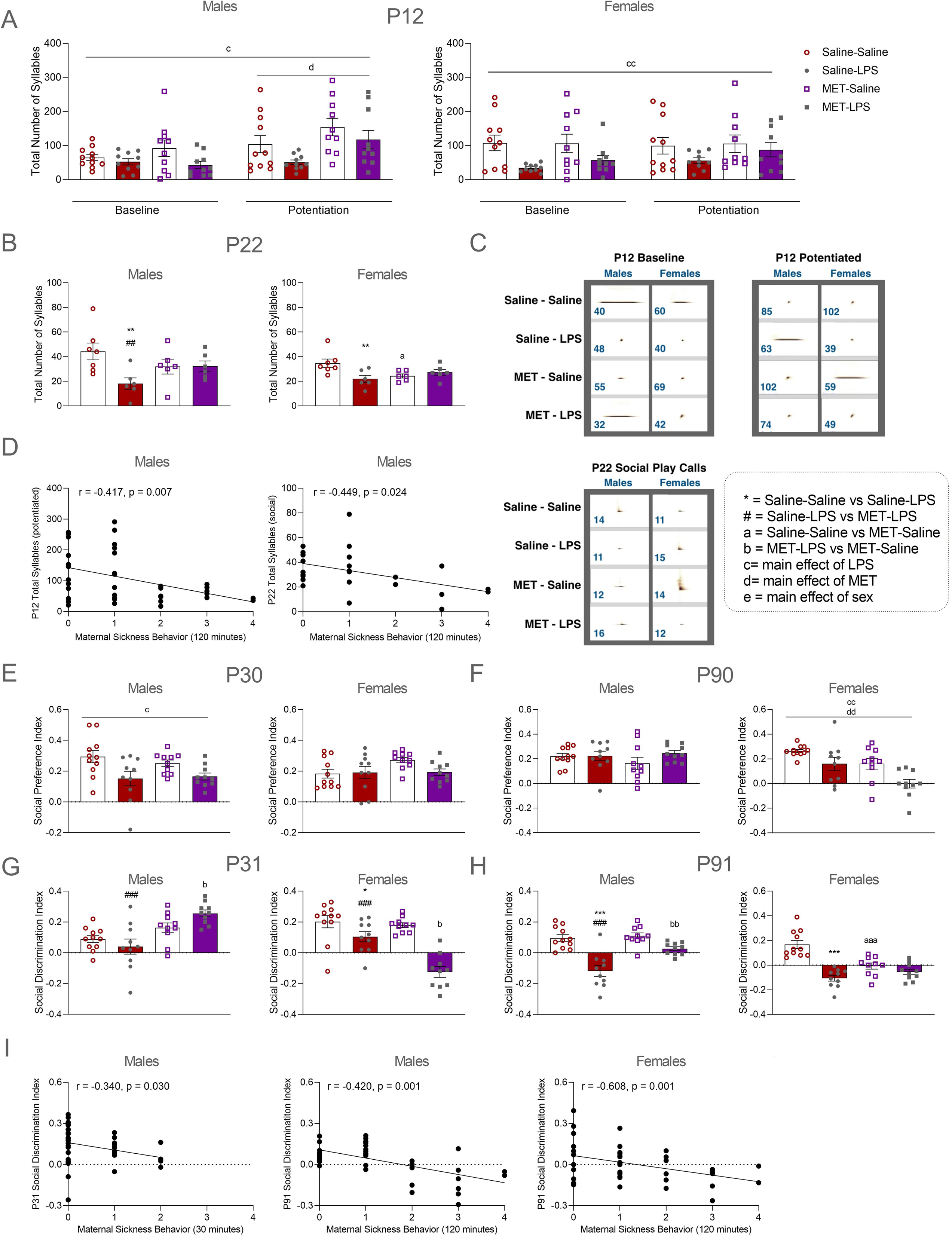
Effect of maternal immune activation (MIA) and metyrapone (MET) on offspring behavioral measures in infancy, adolescence, and adulthood. **A** Neonatal isolation induced ultrasonic vocalizations (USVs) on postnatal day (P)12. MIA reduced the number of syllables produced by males and females during the baseline and potentiation periods. During the potentiation period, MET exposed males emitted a higher total number of syllables compared to unexposed groups. **B** MIA decreased social play calls in males and females on P22 and MET prevented this decrease in male MIA offspring. **C** The top vocalization type, and the number of vocalizations of that type made, for each group during USV tests on P12 and P22. **D** Correlations between MIA induced maternal sickness behavior and total syllables emitted by male offspring on P12 and P22. As the severity of maternal sickness increased, the total number of syllables emitted by male offspring decreased on P12 and P22. **E** Social preference scores were reduced by MIA exposure in male offspring only on P30. **F** On P90, both LPS and MET exposure reduced social preference scores in females only. **G, H** Social discrimination scores were impaired in Saline-LPS males and females on P31 and P91. MET was protective of this effect in males and exaggerated the impairment in female MIA animals across both developmental timepoints. **I** Correlations between MIA induced maternal sickness behavior and social discrimination index. As the severity of maternal sickness increased, social discrimination ability was impaired in adolescent and adult animal offspring. Data are expressed as mean ± SEM. ***p < 0.001, **p < 0.01, *p <0.05., Saline-Saline vs Saline-LPS; ^###^p < 0.001, ^##^p < 0.01, Saline-LPS vs MET-LPS; ^aaa^p < 0.001, Saline-Saline vs MET-Saline; ^cc^p < 0.01, ^c^p < 0.05, main effect of LPS/Saline; ^dd^p < 0.01, ^d^p < 0.05, main effect of MET/Saline. LPS – lipopolysaccharide.

On P30, social preference was reduced following MIA in males but not females (male main effect of LPS: p = 0.05; females: p > 0.05; **Figure 2E**). This observation was reversed on P90 when MIA exposed females (yet not males) had reduced social preference (female main effect of LPS: p = 0.01 and main effect of MET: p = 0.01; males: p > 0.05; **Figure 2F**). Interestingly, P31 social discrimination ability was improved by MET (MET-LPS vs. Saline-LPS; p = 0.001; MET-LPS vs. MET-Saline, p = 0.05; **Figure 2G**). In adulthood, there was a sustained effect of LPS on social discrimination in males (Saline-Saline vs. Saline-LPS: p = 0.001). MET was modestly protective to males at this age (Saline-LPS vs MET-LPS: p = 0.001; **Figure 2H**). Similar to males, there was a sustained effect of LPS on social discrimination in females (Saline-Saline vs. Saline-LPS, p = 0.05; **Figure 2G,H**), however MET exacerbated this deficit in adolescence (MET-LPS vs. Saline-LPS; p = 0.001; **Figure 2G**). While P31 social discrimination in males was only modestly correlated to maternal sickness behaviors 30-minutes post MIA (r = -0.340, p = 0.030; **Figure 2I, Supplemental Results Statistical Table 2**), adult social discrimination was significantly and negatively correlated to the severity of maternal sickness across multiple time points in both sexes (p < 0.05; **Figure 2I; Supplemental Results Statistical Table 2)**. The percent time spent in the center of the open field was only reduced in males by MET on P30 (main effect of MET: p = 0.01; **Supplemental Results Figure 3A**); female Saline-LPS animals spent more time in the center compared to Saline-Saline animals on P90 (p = 0.001; **Supplemental Results Figure 3B**). The total distance traveled was not affected in any group across development (p> 0.05; **Supplemental Results Figure 3C,D**).

### MET reversed transcriptional effects of MIA in the ventral hippocampus of male offspring

RNA-sequencing was performed on ventral hippocampal tissue from adult male and female offspring to characterize transcriptional effects of gestational LPS and MET. The transcriptional effect of LPS exposure and MET exposure (**Supplementary Results Figure 4A,B**) were quite distinct in males and females; therefore, subsequent transcriptional analyses were performed separately in males and females. In males, 290 genes were DE by LPS exposure (i.e., Saline-Saline vs Saline-LPS; **Figure 3A**) and 571 genes were DE by MET exposure (i.e., Saline-LPS vs. MET-LPS; **Figure 3B**). In females, 1,322 genes were DE by LPS (**Figure 4A**), and 513 genes were DE by MET exposure (**Figure 4B**). Notably, threshold-free RRHO analysis revealed that the transcriptional alterations associated with LPS were largely reversed by MET in males (**Figure 3C**) and to a lesser extent in females (**Figure 4C**). Heatmap clustering primarily revealed upregulation of genes belonging to GABA/ glutamate (**Figure 3D**), and monoamine pathways (**Figure 3E-F**) related to offspring development in LPS-exposed males (**Figure 3C**). Many of these LPS-induced alterations were reversed by MET. For example, *Ephb2* (GABA/glutamate pathways), *Bhlhe22*, *Hapln4*, *Camk4* (dopamine pathways), and *Tph2*, *Slc29a4* (serotonin pathways; p < 0.05 and FC > 1.3; **Figure 3D-F**). Similarly, LPS induced alterations in many genes related to GABA, glutamate, and monoamine pathways in females, particularly among genes related to serotonin (**Figure 4D-F**). Although MET altered gene expression in these pathways in females, there were no strong patterns of up or downregulation and no effects were in opposition to the expression changes in LPS-exposed offspring (**Figure 4D-F**). Pathways related to HPA function, immune activation, and offspring development were also identified (**Supplemental Results Figures 5-6**). A summary of gene expression patterns from the other heatmap clusters is available in **Supplemental Results Table 3**. Among other pathways, LPS exposure had a more robust effect in female compared to male offspring, particularly among inflammation-, microglia-, parvalbumin-, and oxytocin-related genes (**Supplemental Results Figures 5-6**). MET generally upregulated genes related to inflammation in males and females. In males, MET led to some upregulation of genes relevant to epigenetics and GR binding. while in females, MET was associated with downregulation of microglial genes and a mix of up and down regulation of genes for parvalbumin and oxytocin. For more information on the genes included in heatmap clustering, see **Supplemental Results Tables 4 and 5.** There were no effects of gestational LPS on GR or CRFR1 protein expression in adult male or female offspring (p > 0.05; **Supplemental Results Figure 7A,B**).

**Figure 3.**
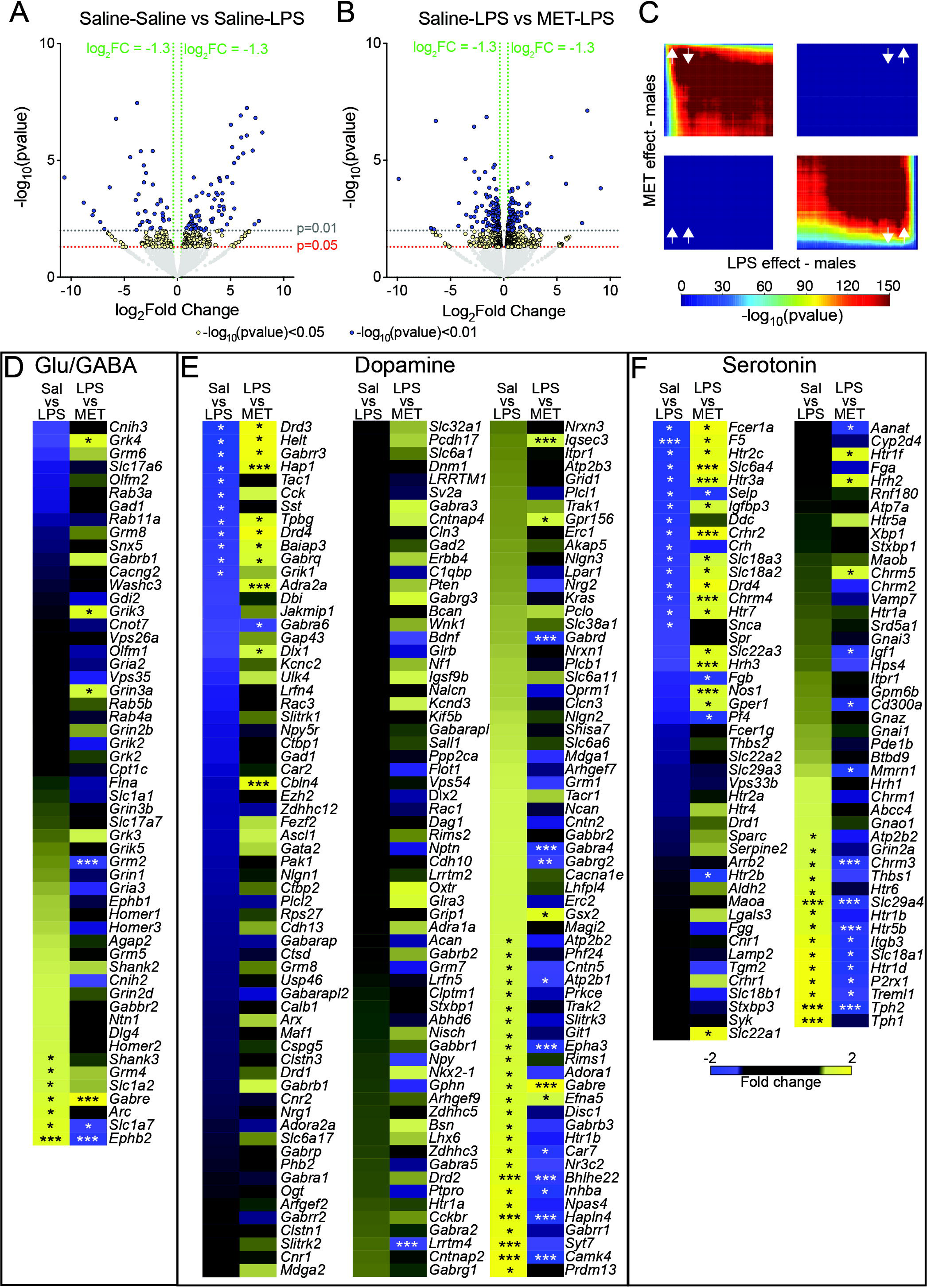
Transcriptomic analyses of ventral hippocampus tissue samples collected from adult male offspring. **A** Volcano plot depicting the distribution of 19,172 genes based on log2 fold change and –log10 p values in saline-saline compared to saline-LPS-exposed offspring. **B** Volcano plot depicting the distribution of 19,194 genes in saline-LPS compared to MET-LPS exposed offspring. Each dot represents a gene, with colors indicating the magnitude of significance (yellow = p < .05, FC > 1.3; blue = p < .01, FC > 1.3). **C** Threshold-free RRHO analysis indicates that the transcriptional alterations induced by LPS are largely reversed by MET. Heatmaps of glutamate (Glu)/GABA (**D**), dopamine (**E**), and serotonin (**F**) related genes, with effects represented as fold change (FC) between saline-saline and saline-LPS-exposed males, and between saline-LPS and MET-LPS-exposed males. Data are expressed as * FC > 1.3, ** p < 0.05, or *** p < 0.05 and FC > 1.3.

**Figure 4.**
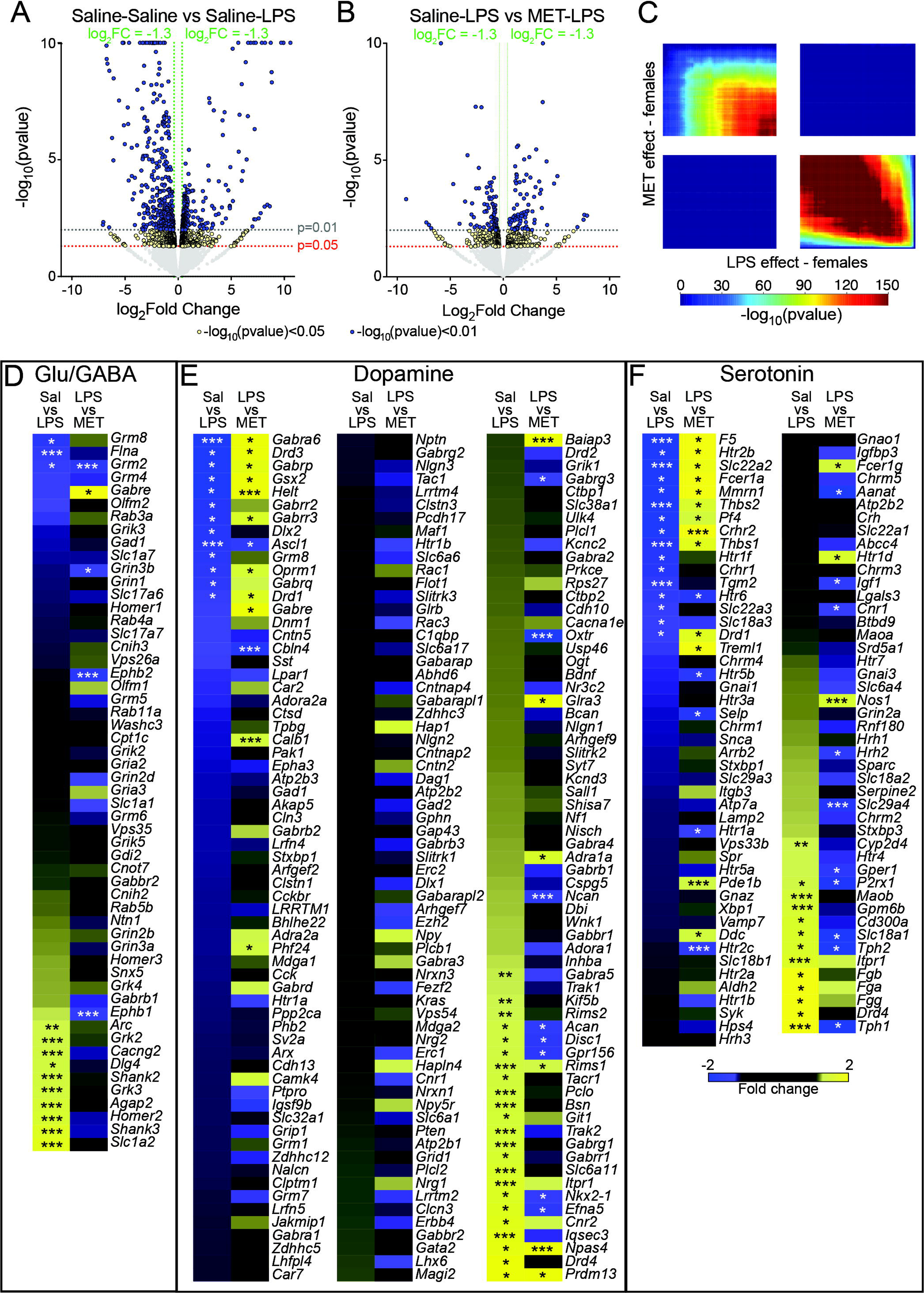
Transcriptomic analyses of ventral hippocampus tissue samples collected from adult female offspring. **A** Volcano plot depicting the distribution of 19,206 genes based on log2 fold change and –log10 p values in saline-saline compared to saline-LPS exposed offspring. **B** Volcano plot depicting the distribution of 19,184 genes in saline-LPS-compared to MET-LPS exposed offspring. Each dot represents a gene, with colors indicating the magnitude of significance (yellow = p < .05, FC > 1.3; blue = p < .01, FC > 1.3). **C** Threshold-free RRHO analysis indicates that the transcriptional alterations induced by LPS are largely reversed by MET. Heatmaps of glutamate (Glu)/GABA (**D**), dopamine (**E**), and serotonin (**F**) related genes, with effects represented as FC between saline-saline and saline-LPS exposed females, and between saline-LPS and MET-LPS-exposed females. Data are expressed as * FC > 1.3, ** p < 0.05, or *** p < 0.05 and FC > 1.3.

## Discussion

Here we show that the 11β-hydroxylase antagonist MET protected against MIA-induced social deficits in a sex-dependent manner. While male MIA offspring appeared to benefit from this intervention, female animals in contrast demonstrated exaggerated social impairments when MET was used in conjunction with MIA. A summary of the main findings and proposed mechanisms in this study can be found in **Figure 5**. Given that MET is used to treat hypercortisolism in pregnant (**Azzola et al., 2020; Kersten et al., 2020; Bass et al., 2021**), lactating (**Duke et al., 2019**) and pediatric populations (**Lourenço et al., 2015; de Mingo et al., 2020**), further assessments of sex-dependent clinical outcomes using longitudinal behavioral studies may be warranted. With regard to MIA, in males, this 11β-hydroxylase antagonist shows some promise to offset the consequences of severe illness during pregnancy. Additional studies in females are needed to determine if adjustments to dose or timing of metyrapone could be beneficial, or if the underlying mechanisms differ enough that alternative interventions must be sought.

**Figure 5.**
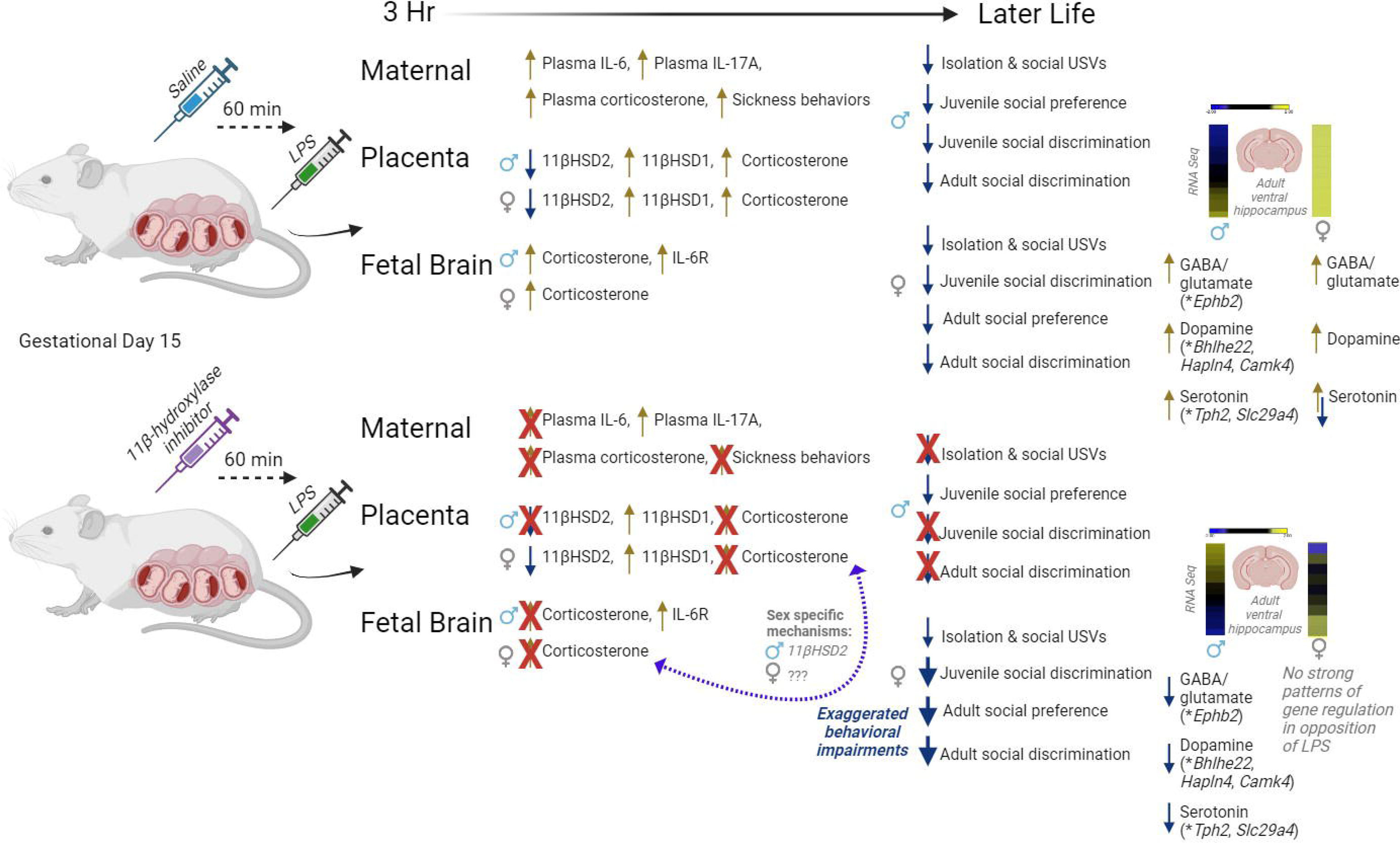
Summary of findings and proposed mechanisms. Lipopolysaccharide (LPS) was administered to pregnant rat dams on gestational day 15 to induce maternal immune activation (MIA). Three hours after injection, corticosterone was elevated in maternal plasma, male and female placentas, and fetal brains. Treating dams with the 11β-hydroxylase inhibitor metyrapone (MET) prior to LPS exposure attenuated maternal sickness behaviors and prevented increases in maternal, placental, and fetal corticosterone concentrations. MET did not affect LPS-induced increases in placental enzyme 11βHSD1 in males or females. However, MET prevented LPS-induced decreases in 11βHSD2 in male placentas only, suggesting that more corticosterone was inactivated in male offspring compared to female offspring. While maternal MET exposure was protective against impairments in ultrasonic vocalizations and social behaviors in males, MET did not protect against behavioral deficits in females and led to further impairments. Prenatal MET exposure also prevented LPS-induced upregulation of several genes related to serotonin, dopamine, GABA, and glutamate in the ventral hippocampus of male, but not female, offspring. These pathways are significant because glucocorticoids are known to influence monoamine pathways and excitatory/inhibitory signaling in the brain. Furthermore, affected genes such as *Ephb2, Camk4,* and *Tph2m* have been implicated in cognitive function and neurodevelopmental disorders. In summary, the HPA axis is activated in conjunction with the immune system to produce MIA-induced deficits in offspring social behaviors across the lifetime. While maternal MET exposure was unable to prevent behavioral impairments in female offspring, MET showed promise as an intervention in males by counteracting MIA effects on 11βHSD2, social behavior and transcriptional changes in the ventral hippocampus.

Like other prenatal stressors (**Haq et al, 2021**), MIA activated the HPA axis by 3hrs post challenge, as evidenced by elevated maternal plasma corticosterone, which subsequently increased corticosterone concentrations in both male and female placentas and fetal brains. It is unclear if the fetally measured corticosterone comes from the dam, from activity of the fetal HPA axis, or from some combination of the two. The fetal HPA axis is immature during pregnancy (**Gunn et al., 2013**), however endogenous secretions of corticosterone have been recorded as early as day 16 of gestation (**Boudouresque et al., 1998**). It is possible that sex differences in behavior arise from differences in endogenous HPA activation, that could result in long-lasting influences on HPA activation and function (**Gunn et al., 2013**). While it is clear that offspring are exposed to corticosterone, further investigation is needed to understand how corticosterone, and 11β-hydroxylase inhibition, may influence the developing HPA axis differently across sex.

In human studies, prenatal stress has been linked to changes in placental enzyme activity and cortisol levels (**Conradt et al., 2013**). Specifically, increased methylation of the gene encoding for 11βHSD2 resulted in silencing of the gene and decreased placental inactivation of cortisol, which is associated with disrupted neurodevelopmental outcomes (**Marsit et al., 2012**). Here, MET-exposure protected against MIA-induced decreases in placental 11βHSD2 in males only, suggesting that less corticosterone was inactivated in female placentas. This further suggests that the combined effects of MIA and MET influence placental corticosterone activity differently between males and females, potentially leading to long-term HPA dysfunction and subsequent behavioral disruption among females.

While MET inhibited maternal sickness behaviors and plasma IL-6 responses in our MIA dams, others have shown sustained elevations in IL-6 following treatment with this 11β-hydroxylase inhibitor (**Alexander & Fewell, 2011; Cui et al., 2011**). Differences in plasma IL-6 levels could be due to variation in the timing of sample collection (**Cui et al., 2011**) or LPS serotype or dose (**Alexander & Fewell, 2011**). While **Hsiao and Patterson (2011**) showed that offspring exposed to gestational IL-6 alone developed schizophrenia- and autism-like behavioral changes consistent with those present in offspring exposed to MIA via the viral mimic poly(I:C), **Bermick and colleagues (2023)** showed that maternal IL-6 does not directly cross into fetal circulation but stimulates an independent fetal IL-6 response. Among offspring in our study, IL-6R expression was greater in MIA-exposed male fetal brains. Notably, this expression remained elevated in MET-LPS offspring, suggesting that an immune response was present, yet protection was still conferred against adverse behavioral outcomes. Together, these findings suggest that cytokines and corticosterone play concerted roles in the detrimental effects of MIA again highlighting the sex-specific nature of these effects.

Atypical ultrasonic vocalizations (USVs) have been associated with symptoms of neurodevelopment and neuropsychiatric disorders in rodents (**Scattoni et al., 2009**). **Malkova and colleagues (2012)** found that male pups prenatally exposed to MIA emitted fewer USVs when separated from their litters and mothers, compared to unexposed pups. Due to the reduced number of harmonic syllables used in these USVs, the authors postulated that the pups exhibited an attenuated emotional response when separated from their dams. Emission of fewer calls when separated from a caretaker might diminish the chance of being found and retrieved. However, it could also be adaptive in that predators would also be unable to find the separated pup (**Zhao et al., 2021**). In another study of USVs recorded during social play, fewer calls were made by adolescent male and female pups that had been exposed to MIA (**Gzielo et al., 2021**). The reduced social calls may result from deficits in sensory information processing, like those observed among individuals with autism. Here, MIA offspring demonstrated altered communication by producing fewer syllables during maternal isolation and social experiences, compared to unexposed pups. MET blocked decreases in P12 potentiated isolation and P22 social play calls, only among male offspring. This finding demonstrates that impairments in USVs are, at least in part, driven by HPA dysregulation.

MIA is associated with sex-dependent social and cognitive impairments that appear later in life (**Malkova et al., 2012; Meyer, 2014; Haida et al., 2019**). Here, preference for a novel rat compared to a novel object at P30 was decreased in MIA-exposed males, yet not females. By adulthood, social preference was recovered in males and was reduced in females exposed to MIA and/or MET. This may indicate that too much, or too little, prenatal corticosterone exposure negatively impacts future social behaviors. In the social discrimination task, rats are presented with a novel animal and a previously encountered animal and must draw on memory to inform their social interactions. In adolescence and adulthood, both MIA males and females displayed deficits in the task, indicating that some aspects of social behavior are affected in both sexes, as shown previously (**Núñez Estevez et al., 2020; Bitanihirwe et al., 2010**). While MET blocked this effect in males, it exacerbated the effect in females, further suggesting that MET modulates the influence of MIA in a sex dependent manner This is not unusual, as the effects of pharmaceutical drugs are commonly found to vary across sexes (**Zucker & Prendergast, 2020; Yoon et al., 2021; Porreca & Dodick, 2023**). Future work is needed to determine if the exaggerated social impairments in female animals can be avoided by adjusting the dose or timing of MET administration and its influence on placental 11βHSD2.

Prenatal exposure to excess glucocorticoids is associated with sex- and region-specific changes in central expression of corticotrophin-releasing factor receptors (CRFRs), downregulation of glucocorticoid receptors (GRs), and long-term dysregulation of the HPA axis (**Zohar & Weinstock, 2011; Liu & Nusslock, 2018**). Downregulated GR expression is indicative of decreased HPA axis negative feedback, which leads to excess corticosterone release and, therefore, a potentiated response to stressors (**Liu & Nusslock, 2018**). Altered CRFR signaling can similarly dysregulate HPA function and subsequent behavior, including increased stress and anxiety, as well as impairments in learning, memory, and social behavior (**Chen et al., 2012; Sukhareva, 2021; Hostetler and Ryabinin, 2013**). Surprisingly, there were no group differences in protein levels of GR or CRFR1 in the ventral hippocampus of male or female offspring, at least in adulthood. Prior work has shown that at P30, relative expression of CRFR1 (CRFR2/CRFR1) mRNA in the hippocampus was reduced and may contribute to impaired social discrimination in males and females exposed to prenatal MIA, compared to animals that were not exposed (**Núñez Estevez et al., 2020**). In the present study, performance on the social discrimination task was similarly impaired in adulthood, suggesting that differences in CRFR1 gene expression that weren’t apparent at the protein level could still be present. RNA-sequencing data revealed changes in CRFR associated genes in MET-exposed male and female adult offspring, pointing to a potential avenue of MET mediated protection. In addition to regulating stress responses in the brain, CRFRs are regulators of monoaminergic signaling (**Magalhaes et al., 2010; Issler et al., 2014; Pleil & Skelly, 2018**). LPS-exposed offspring displayed altered expression of multiple genes associated with serotonin and dopamine pathways, some of which are known to be involved in the etiology of neurodevelopmental disorders (**Garbarino et al., 2019; Ottenhof et al., 2018; Zech et al., 2018; Sałaciak et al., 2021**). Notably, MET exposure reversed many of the LPS-induced gene expression changes in males, but not among females, suggesting that serotonin- and dopamine-related genes could mediate sex differences in the protective effects of MET on social behaviors in offspring. Similarly, MIA exposure in females resulted in upregulation of GABA- and glutamate-related genes, many of which are also implicated in neurodevelopmental deficits (**Fiorentino et al., 2015; Sellgren et al., 2021; Huang et al., 2019**), although MET exposure had no effect on their expression. In contrast, there was only one gene, *EphB2*, that was significantly upregulated in this family among males, and the effect was opposed by MET. This particular gene and its protein have previously been implicated with autism-like social behavioral displays in mice (**Assali et al., 2021; He et al., 2023; Liu et al., 2021; Wu et al., 2020**). In sum, the effect of LPS-exposure is multifaceted, and MET-exposure may protect against social deficits more effectively in male compared to female offspring.

In summary, attenuating corticosterone exposure in pregnant rat dams diminishes the presence of some MIA-induced behavioral outcomes in offspring; these effects are sex dependent and vary across the course of development. Future work is necessary to determine if administration dose or time effects underlie the exaggerated impairments observed in female MIA offspring. MET is already used in pregnant and pediatric populations, so this 11β-hydroxylase inhibitor could be a promising and feasible intervention to attenuate maternal HPA activation during severe gestational illness, at least for males. Long-term behavioral studies designed to investigate potential sex-specific effects of MET exposure in utero and in early childhood may be warranted in clinical settings. Indeed, it is possible that MET administration could be detrimental to female offspring, important information to know given this drug’s clinical use across pregnancy and lactation.

## Supporting information

Supplementary Table. MIA Guidelines Reporting Table

Supplementary Methods, Supplementary Figures and Smaller Supplementary Tables

Supplementary Table. Statistical Reporting of Maternal and Offspring Measures

Supplementary Table. Gene Expression in the Adult Male Ventral Hippocampus

Supplementary Table. Gene Expression in the Adult Female Ventral Hippocampus

## Author Contributions

J.M., L.G., & M.A.S ran the experiments; J.M, M.A.S., M.L.S., & A.C.K. analyzed and interpreted the data; J.M and A.C.K. wrote the manuscript; A.C.K., designed and supervised the study.

## Data availability

RNA-seq data has been deposited to GEO (GSE240604). All other data is available on request.

## Code availability

There is no code associated with this work.

## Figure Captions

**Supplementary Materials.** Supplementary Materials. Extended Supplementary Methods, Supplementary Table 1, Supplemental Figures 1-7, Supplemental Table 3.

**Supplemental Methods Table 1.** Maternal Immune Activation (MIA) reporting guidelines checklist.

**Supplemental Results Figure 1. Effect of maternal immune activation (MIA) and metyrapone (MET) on western blot analyses of receptor and cytokine expression in fetal brain tissue. A** Corticotropin-releasing factor receptor 1 (CRFR1), **B** glucocorticoid receptor (GR), and **C** IL-17A densitometric ratios in male and female fetal brain tissue. CRFR1/ß-Actin, GR/ß-Actin, and IL-17A/ß-Actin ratios were not significantly different between groups 3-hours post challenge. Data are expressed as mean ± SEM. LPS – lipopolysaccharide.

**Supplemental Results Figure 2. Effect of maternal immune activation (MIA) and metyrapone (MET) on mean duration of ultrasonic vocalizations (USVs) in offspring. A** There were no differences between groups in the mean duration (msec) of isolation induced USVs made during the baseline and potentiation periods on postnatal day (P)12. **B** There were also no group differences in the mean duration (msec) of USV social play calls made on P22. Data are expressed as mean ± SEM. LPS – lipopolysaccharide.

**Supplemental Results Figure 3. Effect of maternal immune activation (MIA) and metyrapone (MET) on offspring open field and distance traveled measures. A** MET exposed males spent less time in the center of the open field on P30, compared to unexposed males (main effect of MET/Saline: p = 0.01). **B** On P90, Saline-LPS (p = 0.001) and Saline-MET (p = 0.01) females spent more time in the center of the apparatus than Saline-Saline females. **C, D** Distance travel was not affected in any group on P30 or P90. Data are expressed as mean ± SEM. **p < 0.01, Saline-Saline vs Saline-LPS; ^ddd^p < 0.001, main effect of MET/Saline. LPS – lipopolysaccharide.

**Supplementary Results Figure 4. Maternal immune activation (MIA) and metyrapone (MET) induce sex-specific transcriptional effects in the ventral hippocampus. A** Threshold-free rank-rank hypergeometric overlap (RRHO) analysis revealed distinct transcriptional effects in males and females exposed to lipopolysaccharide (LPS). **B** RRHO revealed distinct transcriptional effects in males and females exposed to MET.

**Supplemental Results Figure 5. Transcriptomic analyses of ventral hippocampus tissue samples collected from adult male offspring. A** Heatmaps of inflammation-, microglia-, LPS binding-, oxytocin-, epigenetics-, prolactin-, parvalbumin-, aldosterone/MR signaling-, glucocorticoid signaling-, GR binding-related genes, with effects represented as fold change (FC) between saline-saline and saline-LPS-exposed males, and between saline-LPS and MET-LPS-exposed males. Data are expressed as * FC > 1.3, ** p < 0.05, or *** p < 0.05 and FC > 1.3.

**Supplemental Results Figure 6. Transcriptomic analyses of ventral hippocampus tissue samples collected from adult female offspring. A** Heatmaps of inflammation-, microglia-, LPS binding-, oxytocin-, epigenetics-, prolactin-, parvalbumin-, aldosterone/MR signaling-, glucocorticoid signaling-, GR binding-related genes, with effects represented as FC between saline-saline and saline-LPS exposed females, and between saline-LPS and MET-LPS-exposed females. Data are expressed as * FC > 1.3, ** p < 0.05, or *** p < 0.05 and FC > 1.3.

**Supplemental Results Figure 7. Effect of maternal immune activation (MIA) and metyrapone (MET) on western blot analysis of receptor expression in the ventral hippocampus of adult offspring.** Ventral hippocampal **A** glucocorticoid receptor (GR) and **B** corticotropin-releasing factor receptor 1 (CRFR1) densitometric ratios for male and female offspring on P94. Neither GR/ß-Actin ratios nor CRFR1/ß-Actin ratios were significantly different between groups. Data are expressed as mean ± SEM. LPS – lipopolysaccharide.

**Supplemental Results Table 1.** Total numbers of animals and litters in each measure.

**Supplemental Results Table 2.** Statistical reporting table of maternal and offspring measures.

**Supplemental Results Table 3.** Summary of gene expression patterns in heatmap clusters

**Supplemental Results Table 4.** Gene expression in the adult male ventral hippocampus

**Supplemental Results Table 5.** Gene expression in the adult female ventral hippocampus

