## Supplementary Methods, Supplementary Figures and Smaller Supplementary Tables for "Sex differences in offspring risk and resilience following 11β-hydroxylase antagonism in a rodent model of maternal immune activation"

Running title: MIA and 11β-hydroxylase antagonism

Julia Martz[1], Micah A. Shelton[2], Laurel Geist[1], Marianne L. Seney[2], Amanda C. Kentner[1]*

[1]School of Arts & Sciences, Health Psychology Program, Massachusetts College of Pharmacy and Health Sciences, Boston Massachusetts, United States 02115.

[2]Department of Psychiatry, University of Pittsburgh, 450 Technology Drive Pittsburgh, PA, 15219.

*Corresponding author:
Amanda Kentner

Office #617-274-3360
Fax # 617-732-2959

### Supplementary Materials

#### Supplementary Materials & Methods

##### Animals and Experimental Overview

All animal procedures were approved by the Massachusetts College of Pharmacy and Health Sciences (MCPHS) Institutional Animal Care and Use Committee and were carried out in line with the Association for Assessment and Accreditation of Laboratory Animal Care. The experimental procedures and groups evaluated in this study are presented in **Figure 1A** and **Supplementary Results Table 1**. **Supplementary Methods Table 1** provides additional methodological details (Kentner et al., 2019). Sprague Dawley rats were acquired from Charles River Laboratories (Male breeders: Wilmington, MA; Maternal and fetal tissue studies: Hollister, CA; Behavioral studies: Raleigh, NC) and housed on a light/dark cycle (0700-1900 light at 20°C) with *ad libitum* access to food and water. After acclimatizing to the animal facility, animals were bred (1 male to 2 females) and pregnancy confirmed by the visualization of spermatozoa in vaginal samples, a sustained diestrus phase, and constant body weight gain.

On the morning of gestational day (G) 15, rat dams were administered either pyrogen-free saline (vehicle) or the 11β-hydroxylase inhibitor metyrapone (MET, 100 mg/kg, i.p). One hour later, dams were treated with either the vehicle pyrogen-free saline, or 200 μg/kg (i.p.) of the inflammatory endotoxin lipopolysaccharide (LPS; Escherichia coli, serotype 026:B6; L-3755) to induce MIA. To validate MIA induction by LPS, sickness behaviors were evaluated (Experiment 1; n = 7-14). Passive observations of dams in their home cages were made on measures of ptosis (droopy eyelids), piloerection (ruffled and unkempt coat appearance), and lethargy. A blind observer scored each of these behaviors on a three-point scale (0 = none, 1 = mild, 2 = severe) as previously described (Connors et al., 2014) at baseline, 30, 60, 90-, 120-, 240-, and 360-minutes post injection. A composite score was then assigned for each time point (adapted from Hayley et al., 2002; Kentner et al., 2007) where: 0 = 'no sickness' indicated by all behaviors being scored as none, or one behavior scored as mild; 1 = two or more mild scores, no severe scores (i.e., mild lethargy and ptosis and/or piloerection); 2 = severe score on one sickness behavior (mild or none scored on all other behaviors); 3 = severe score on two sickness behaviors; 4 = ‘very sick appearance’ as indicated by 3 severe scores (severe ptosis, lethargy, and piloerection). A subset of these dams was euthanized by ketamine/xylazine (150 mg/kg/50 mg/kg, i.p.) 3 hours following the G15 LPS challenge time point (n = 7). Maternal cardiac plasma samples were collected and stored for further validation of MIA by ELISA. Dams were perfused intracardially with a phosphate buffer solution (PBS) and the uterine horn removed. Placentas, whole fetal brains, and bodies were also saved for later processing. The first three conceptuses from either side of the cervical end were selected for evaluation, to limit effects of intrauterine position (Bronson & Bale, 2014). The remaining dams evaluated in the 6-hr sickness behavior observations were euthanized the following day (ketamine/xylazine: 150 mg/kg/50 mg/kg, i.p.).

A second set of dams was used to validate MET by confirming the 11β-hydroxylase inhibitor attenuated the sustained release of maternal plasma corticosterone in response to LPS (Experiment 2; n = 5-7). The tails of these dams were lanced close to the tip and EDTA coated Microvettes® (CB 300 K2E; Sarstedt, Germany) were used to collect tail blood at 60, 90-, 120-, 240-, and 360-minutes post LPS challenge. Samples were kept on ice, centrifuged and plasma aliquots were stored at −80°C until processing.

Following the G15 drug treatments, a third group of dams (Experiment 3; n = 10-11) were permitted to continue their pregnancy until birth (postnatal day (P)1) and their offspring evaluated on several behavioral metrics. On G19, these dams were separated into individual cages (27 × 48 × 20 cm). On P2, the pups were standardized to 10 pups per litter (5 males and 5 females, wherever possible) and stayed with their mothers until weaning on P21. At that point, offspring were placed into clean cages in same-sex pairs where they were maintained throughout the rest of the study. Behavior was evaluated in one male and one female from each litter per measure during the juvenile (P12 and P22) and adolescent (P30 and P31) periods and again as adults (P90-P91).

**Maternal and Fetal Tissue Processing**

To determine the sex of fetuses, genomic DNA was extracted from embryonic tissue (e.g., fetal body) using a DNA extraction kit (Cat. #69504, DNeasy Blood & Tissue Kits, QIAGEN, Valencia, CA, USA), according to manufacturer's instructions. For polymerase chain reaction (PCR), the sequence of the Sry forward primer was: 5′ CGGGATCATGTCAAGCGCCCCATGAATGCATTTATG 3′ and reverse primer 5′ GCGGAATTCACTTTAGCCCTCCGATGAGGCTGATAT 3′. As a control, the sequence of β-actin forward primer 5′ AGCCAT GTACGTAGCCATCC 3′ and reverse primer 5′ TGTGGTGGTGAAGCTGTAGC 3′ was amplified by PCR. The PCR was done by adding 22.5 µL of DNA PCR SuperMix (Cat. #10572014, Thermo Fisher) to a PCR tube, then adding 0.5 µL of SRY forward primer and 0.5 µL of SRY reverse primer. 0.5 µL of β-actin forward primer (1:10 dilution) and 0.5 µL of β-actin reverse primer was added to the PCR tube along with 1 µL of DNA. The PCR conditions were preheating at 105 °C, then setting initial denature to 95 °C for 2 min. The PCR machine was set to 35 cycles, step 1: 95 °C (1 min), step 2: 52 °C (1 min) step 3: 72 °C (1 min), final extension: 72 °C (5 min) and a final hold at 10 °C (protocol adapted from Miyajima et al., 2009; Núñez Estevez et al., 2020). 8 µL of the PCR product was mixed with 2 µL of 6x DNA dye (Cat. #R1151, Thermo Fisher) and loaded onto a 2% agarose gel. Sex was determined by the presence of a Sry band as visualized using an Accuris™ SmartDoc™ imager (Cat. #Z742591-1EA, Millipore Sigma; **Figure 1A)**.

Corticosterone levels of maternal plasma, fetal brains, and placenta were measured in duplicate by ELISA according to the small sample assay protocol of the standard testing kit (Cat. #ADI-900-097, Enzo Life Sciences, Farmingdale, NY). The minimum detectable concentration was 26.99 pg/ml, and the intra- and inter-assay coefficients of variation were 6.6% and 7.8%, respectively.

Maternal concentrations of interleukin (IL)-6 and IL-17A were also measured in duplicate by ELISA (Cat. #ab234570: IL-6 and Cat. #ab119536: IL-17A, Abcam, Waltham, MA). The minimum detectable concentration was 46 pg/ml and 1 pg/ml for IL-6 and IL-17A respectively. The intra-assay coefficients were 2.5% (IL-6) and 8.5% (IL-17A) and inter-assay coefficients were 3.8% (IL-6) and 7.6% (IL-17A).

Placental levels of 11β-hydroxysteroid dehydrogenase (11βHSD1) 1 and 11βHSD2 were similarly measured by ELISA (Cat. # MBS3807536: 11βHSD1 and Cat. # MBS755288: 11βHSD2, MyBioSource, San Diego, CA). The minimum detectable concentration was 0.1 U/L and 0.1 ng/ml for 11βHSD1 and 11βHSD2 respectively. The intra-assay and inter-assay coefficients were <15% (11βHSD1) and <10% (11βHSD2). Fetal brains were further evaluated by Westerns for corticotropin releasing factor receptor 1 (CRFR1), glucocorticoid receptor (GR), IL-6 receptor (IL-6R), and IL-17A (described below).

**Behavioral Analyses**

On P4, a short maternal retrieval test took place. Dams were removed from their litter and all the offspring were pseudorandomly dispersed across the home cage (as previously described by Spencer et al., 2006). The latency (seconds) for dams to retrieve the first pup and the whole litter was recorded. For the remaining behavioral assessments, only one male and one female from each litter were evaluated on each measure (**see Supplemental Results Table 1**).

On P12, pups were individually removed from their home cage and placed into a small cage lined with Nestlets®. To measure maternal potentiation of ultrasonic vocalizations (USVs), the small cage was placed into a sound attenuation chamber and neonatal isolation calls were recorded from an ultrasonic microphone (UltraSoundGate 116–200), located 20 cm from the floor. After briefly reuniting with their dams (for 5 minutes), pups were again removed and tested as above (potentiation period; adapted from Bodi et al., 2016; Kentner et al., 2018). On P22, offspring were introduced to a novel conspecific of the same treatment group and recorded for 5 minutes (as above; Kentner et al., 2018). The USV microphone was connected to a PC and the software program Audacity was used to record vocalizations at 192 kHz, capturing a range of USVs. USVs were analyzed by MUPET 2.0 software (Van Segbroeck et al., 2017; http://sail.usc.edu/mupet/) to determine the total number of syllables emitted and average call duration (msec) from each vocalization recording. Repertoires showing the top syllable types and number emitted by each group during the P12 potentiation and P22 play periods were also generated (MUPET 2.0 software) in order to present a descriptive depiction of USV signature differences (Kentner et al., 2018).

Open field, social preference, and social discrimination tests were each evaluated in adolescent and adult animals as previously described (see Yan & Kentner, 2017; Connors et al., 2014; Zhao et al., 2021a; Núñez Estevez et al., 2020). For open field, distance travelled (m) and time (seconds) spent in each of the perimeter and center of the field was analyzed using automated behavioral monitoring software (Any-maze, Wood Dale, IL) and data presented as a percentage of time spent in the center of the open field [(total time in center/total time in perimeter + total time in center) × 100] as an index of anxiety-like behavior.

Following the open field habituation period, two cleaned wire containment cups were placed on each end of the arena for a five-minute social preference test. One cup held a novel untreated rat of the same sex, age, size, and strain and the other cup held a novel object. Location of novel rats (n = 10) and objects was interchanged between trials. A rat was actively investigating when its nose was directed within 2 cm of a containment cup, or it was touching the cup. For each animal, a social preference index was calculated by the equation ([time spent with the rat] / [time spent with the inanimate object + time spent with the rat]) − 0.5 (Scarborough et al.,

2020). Twenty-four hours later, animals were evaluated on the social discrimination test where one containment cup held the same (now familiar) rat from the day before, and the other cup held a novel animal of the same sex//age/size/strain. The social discrimination index was calculated as ([time spent with the novel rat] / [time spent with the familiar rat + time spent with the novel rat]) − 0.5.

**Western Blot Analyses**

On P94, offspring were anesthetized with a mixture of ketamine/xylazine (150 mg/kg /50 mg/kg, i.p.). Animals were perfused intracardially with PBS, brains removed, placed over ice and the ventral hippocampus dissected. All samples were immediately frozen on dry ice and stored at −80 °C until analysis. To measure GR and CRFR1 expression in adult brain or GR, CRFR1, IL17A, and IL-6 receptor (R) in fetal brain using western blotting, 30 μg of protein was loaded into each well of MiniProtean® gels (Bio Rad Laboratories, Cat. #4568101). Gels were transferred onto nitrocellulose membranes (Bio Rad Laboratories, Cat. #1620147) and blocked in 5% nonfat milk with TBS + 0.05% Tween 20 (TBST) for 1 hour at room temperature (IL-6R, IL-17A, β-actin) or 10% nonfat milk with TBST overnight at 4°C (GR, CRFR1). Membranes were washed with TBST 3x10 minutes and incubated in a 1:1000 dilution of primary antibody in TBS (Thermo Fisher Scientific Cat. #PA5-18801, CRFR1; Cat. #MA1-510, GR for 1.5 hours at room temperature; or Cat. #700480, IL-6R; Cat. #PA5-46947, IL-17A overnight at 4°C; or Cat. #MA515739, β-actin for 1 hour at room temperature). Next, membranes were washed with TBST 3x5 minutes and incubated in a 1:1000 dilution of HRP conjugated secondary antibody (Thermo Fisher Scientific, Cat. #61-0220, GR; Cat. #31402, IL-17A), made in 1% nonfat milk with TBS for 1 hour at room temperature. Otherwise, membranes were washed and incubated in an HRP-conjugated secondary antibody in a 1:500 dilution (Thermo Fisher Scientific, Cat. #32460, IL-6R and IL-17A), made in 1% nonfat milk with TBS, for 1 hour at room temperature. After probing each target with their corresponding secondary antibody, membranes were washed, exposed to chemiluminescent substrate (Thermo Fisher Scientific, Cat. #34580) for 5 minutes and scanned with a LI-COR C-DiGit Scanner (Model # 3600). Densitometry measures were used to obtain a ratio of Antibody target/β-actin in order to quantify differences between groups.

**RNA Sequencing**

Adult ventral hippocampal tissue samples (n=3/group) were homogenized using the Bead Ruptor 12 (Omni International, Kennesaw, GA, USA) and RNA was extracted using RNeasy Plus Micro kits (Qiagen, Valencia, CA, USA). RNA concentration was determined using an Invitrogen Qubit 4 Fluorometer (Thermo Fisher Scientific Inc., Waltham, USA) and RNA integrity was measured with the RNA Nano 6000 Assay Kit of the Agilent Bioanalyzer 2100 system (Agilent Technologies, CA, USA). All RIN values were measured >7.9, indicating excellent RNA quality. RNA libraries were prepared and sequenced at Azenta Life Sciences (South Plainfield, NJ, USA). A NEBNext Ultra II RNA Library Prep Kit for Illumina (NEB, Ipswich, MA, USA) was used to construct RNA sequencing libraries. The manufacturer's instructions were used to enrich mRNAs with Oligod(T) beads, fragment the enriched mRNAs for 15 minutes at 94 °C, and then synthesize first strand and second strand cDNA. End repair and adenylation at the 3' end was performed prior to ligation of universal adapters to the cDNA fragments. Following index addition, a limited cycle PCR was used for library enrichment. An Agilent TapeStation (Agilent Technologies, Palo Alto, CA, USA) system was used to run the sequencing library. Quantification was conducted with a Qubit 2.0 Fluorometer (Invitrogen, Carlsbad, CA) and quantitative PCR (KAPA Biosystems, Wilmington, MA, USA). The sequencing libraries were then clustered and loaded onto the NovaSeq 6000 sequencing system (Illumina, San Diego, CA, USA). The samples were sequenced according to the manufacturer's instructions using a 2x150bp Paired End (PE) configuration. Illumina Control software was used to perform image analysis and base calling. The raw binary base call (.bcl) files generated by the sequencer were converted into fastq files and de-multiplexed using bcl2fastq 2.17 software from Illumina. For index sequence identification, a single mismatch was allowed.

**Statistical Analyses**

Friedman's non-parametric version of the one-way repeated measures ANOVA was used to evaluate maternal sickness behavior. Kruskal-Wallis was used for follow-up tests. Fetal tissues were adequately powered to detect sex-differences, so three-way ANOVAs (Sex x G15 MET/Saline x G15 LPS/Saline) were utilized with litter as a covariate. Two-way ANOVAs were used as appropriate for all other measures (G15 MET/Saline x G15 LPS/Saline) unless the data were skewed (Shapiro-Wilks) then non-parametric Kruskal-Wallis tests were employed (see Green & Salkind, 2005). Because behavioral and adult tissue datasets were not powered to evaluate sex-differences directly, combined with the fact that we were interested in sex-specific effects, male and female animals were evaluated separately (Ordoñes Sanchez et al., 2021). Partial eta-squared ($n_p^2$) is reported as an index of effect size for ANOVAs (Miles & Shevlin, 2001). Pearson correlations were utilized to determine associations for prenatal and postnatal measures. All data are expressed as mean ± SEM.

For RNA sequencing, we used CLC Genomics Workbench v.23.0.2 software (QIAGEN Digital Insights, Aarhus, Denmark) to perform differential expression analysis. Because we were not powered to assess the effect of sex and to coincide with our behavioral analysis approach, males and females were analyzed separately. Our approach was to identify genes in which the effect of LPS was reversed by MET: 1) We first probed for effects of LPS by comparing expression of Saline-LPS animals to the Saline-Saline group; 2) We next compared expression in MET-LPS animals to the Saline-LPS group. Putting these differential expression analyses together, we could determine which genes were altered by LPS for which the effect was reversed by MET. Volcano plots for each brain region were constructed using the log2 fold and -log10 p-value change of expression in GraphPad Prism v.9.5.1. In order to maintain the same size axes across all plots, the -log10 p-value was capped at 10 for all genes. The actual p-value for all genes can be found in the **Supplemental Results Tables 4 and 5**. We were particularly focused on pathways associated with dopamine, serotonin, glutamate, GABA, inflammation, microglia, LPS binding, oxytocin, epigenetics, prolactin, parvalbumin, aldosterone/MR signaling, glucocorticoid signaling, and GR binding; we report patterns of gene expression changes for these pathways using heatmap visualizations of the effects of LPS and MET administration for each sex. Heatmaps representing our expression data were created using the free browser program Morpheus (https://software.broadinstitute.org/morpheus). Genes with $p < 0.05$ and fold change > 1.3 were considered differentially expressed. RRHO2 was used as a threshold-free approach to determine the extent of concordance/discordance between transcriptomic datasets (Cahill et al., 2018).

### Supplementary Results

**Supplemental Results Table 1.** Total numbers of animals and litters in each measure

| Treatment | Number of Animals in Maternal Sickness Behavior Observations (Exp. 1) | Number of Animals in Maternal LPS Validation ELISAs (Exp. 1) | Number of Litters[a] in Fetal ELISA Measures (Exp. 1) | Number of Animals in MET Validation Corticosterone ELISA (Exp. 2) | Number of Offspring[b] Included in P12 Behavior Analyses (Exp. 3) | Number of Offspring[b] Included in P22 Behavior Analyses (Exp. 3) | Number of Offspring[b] Included in P30-31 and P90-91 Behavior Analyses (Exp. 3) | Number of Offspring[b] Included in P94 Ventral Hippocampal Analyses (Exp. 3) |
|---|---|---|---|---|---|---|---|---|
| Saline-Saline | 30 min to 120 min: 14 dams<br><br>240 to 360 min: 7 dams | 3 hrs: 7 dams | Males<br>3 hrs: 7<br><br>Females<br>3 hrs: 7 | 60 to 360 min: 5 dams | Male: 11<br>Female: 11 | Male: 7<br>Female: 7 | Male: 11<br>Female: 11 | Males:<br>Westerns: 6-7<br>RNASeq: 3<br><br>Females:<br>Westerns: 7<br>RNASeq: 3 |
| Saline-LPS | 30 min to 120 min: 14 dams<br><br>240 to 360 min: 7 dams | 3 hrs: 7 dams | Males<br>3 hrs: 7<br>24 hrs: 7<br><br>Females<br>3 hrs: 7 | 60 to 360 min: 7 dams | Male: 10<br>Female: 10 | Male: 6<br>Female: 6 | Male: 11<br>Female: 11 | Males:<br>Westerns: 6-7<br>RNASeq: 3<br><br>Females:<br>Westerns:7<br>RNASeq: 3 |
| MET-Saline | 30 min to 120 min: 14 dams<br><br>240 to 360 min: 7 dams | 3 hrs: 7 dams | Males<br>3 hrs: 7<br><br>Females<br>3 hrs: 7 | 60 to 360 min: 5 dams | Male: 10<br>Female: 10 | Male: 6<br>Female: 6 | Male: 11<br>Female: 11 | Males:<br>Westerns: 6<br>RNASeq: 3<br><br>Females:<br>Westerns: 7<br>RNASeq: 3 |
| MET-LPS | 30 min to 120 min: 14 dams<br><br>240 to 360 min: 7 dams | 3 hrs: 7 dams | Males<br>3 hrs: 7<br>24 hrs: 7<br><br>Females<br>3 hrs: 7 | 60 to 360 min: 6 dams | Male: 10<br>Female: 10 | Male: 6<br>Female: 6 | Male: 11<br>Female: 11 | Males:<br>Westerns: 6<br>RNASeq: 3<br><br>Females:<br>Westerns: 7<br>RNASeq: 3 |

P: Postnatal day, LPS: lipopolysaccharide; MET: Metyrapone
[a]Only fetal tissue from one male and one female were evaluated from each litter.
[b]Only one male and one female pup were evaluated from each litter.

**Supplemental Results Figure 1. Effect of maternal immune activation (MIA) and metyrapone (MET) on western blot analyses of receptor and cytokine expression in fetal brain tissue. A** Corticotropin-releasing factor receptor 1 (CRFR1), **B** glucocorticoid receptor (GR), and **C** IL-17A densitometric ratios in male and female fetal brain tissue. CRFR1/ß-Actin, GR/ß-Actin, and IL-17A/ß-Actin ratios were not significantly different between groups 3-hours post challenge. Data are expressed as mean ± SEM. LPS – lipopolysaccharide.

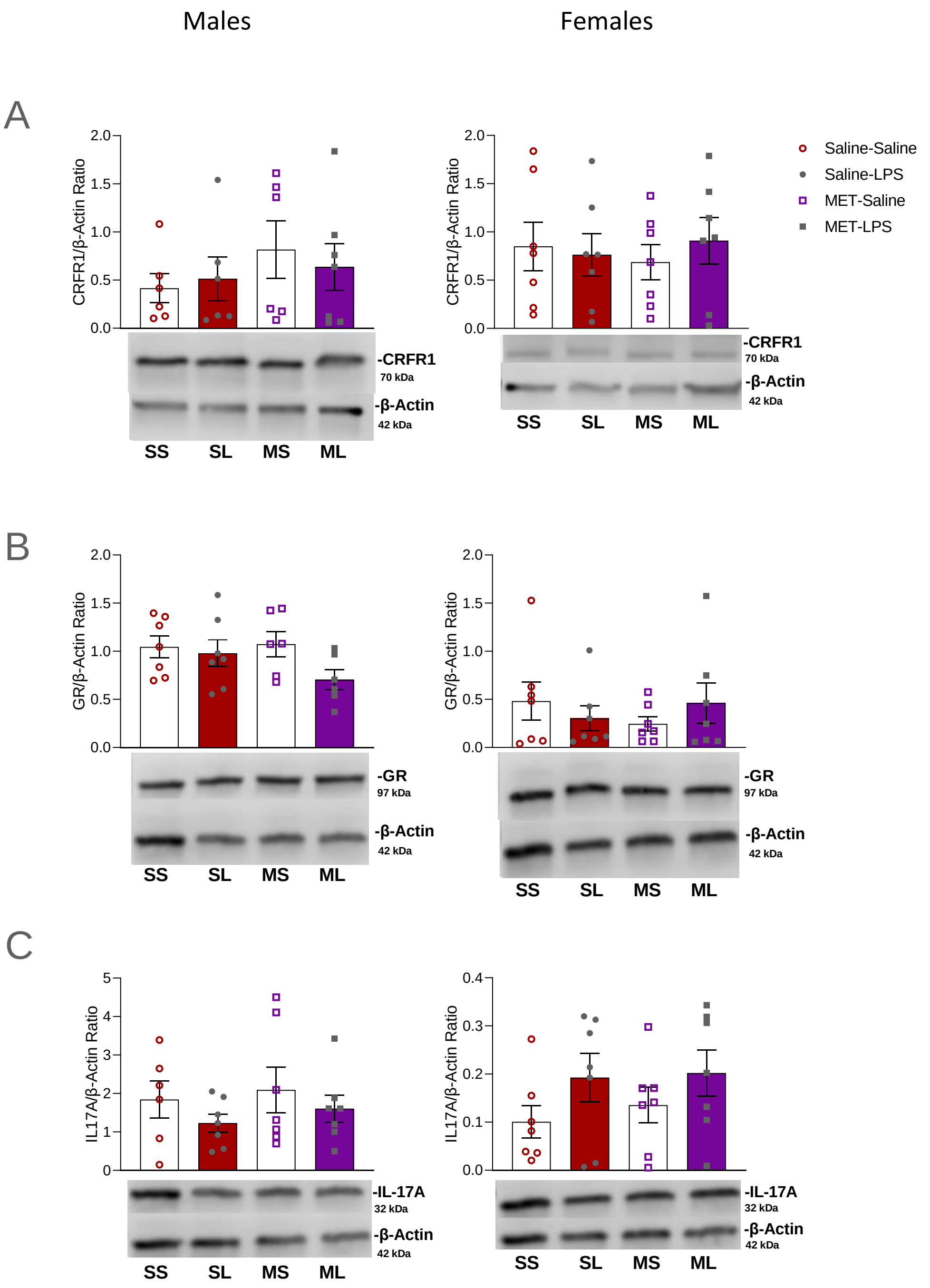

**Supplemental Results Figure 2. Effect of maternal immune activation (MIA) and metyrapone (MET) on mean duration of ultrasonic vocalizations (USVs) in offspring. A** There were no differences between groups in the mean duration (msec) of isolation induced USVs made during the baseline and potentiation periods on postnatal day (P)12. **B** There were also no group differences in the mean duration (msec) of USV social play calls made on P22. Data are expressed as mean ± SEM. LPS – lipopolysaccharide.

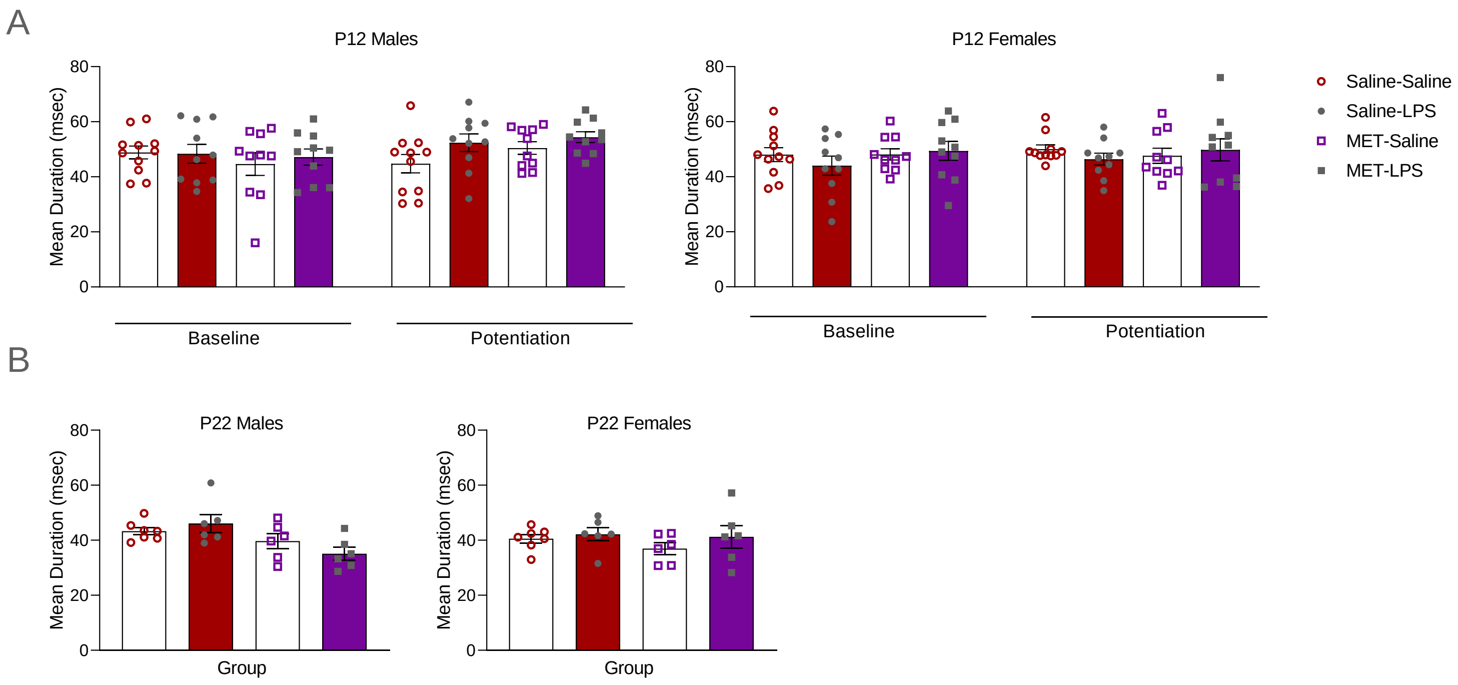

**Supplemental Results Figure 3. Effect of maternal immune activation (MIA) and metyrapone (MET) on offspring open field and distance traveled measures. A** MET exposed males spent less time in the center of the open field on P30, compared to unexposed males (main effect of MET/Saline: $p = 0.01$). **B** On P90, Saline-LPS ($p = 0.001$) and Saline-MET ($p = 0.01$) females spent more time in the center of the apparatus than Saline-Saline females. **C, D** Distance travel was not affected in any group on P30 or P90. Data are expressed as mean ± SEM. $^{**}p < 0.01$, Saline-Saline vs Saline-LPS; $^{ddd}p < 0.001$, main effect of MET/Saline. LPS – lipopolysaccharide.

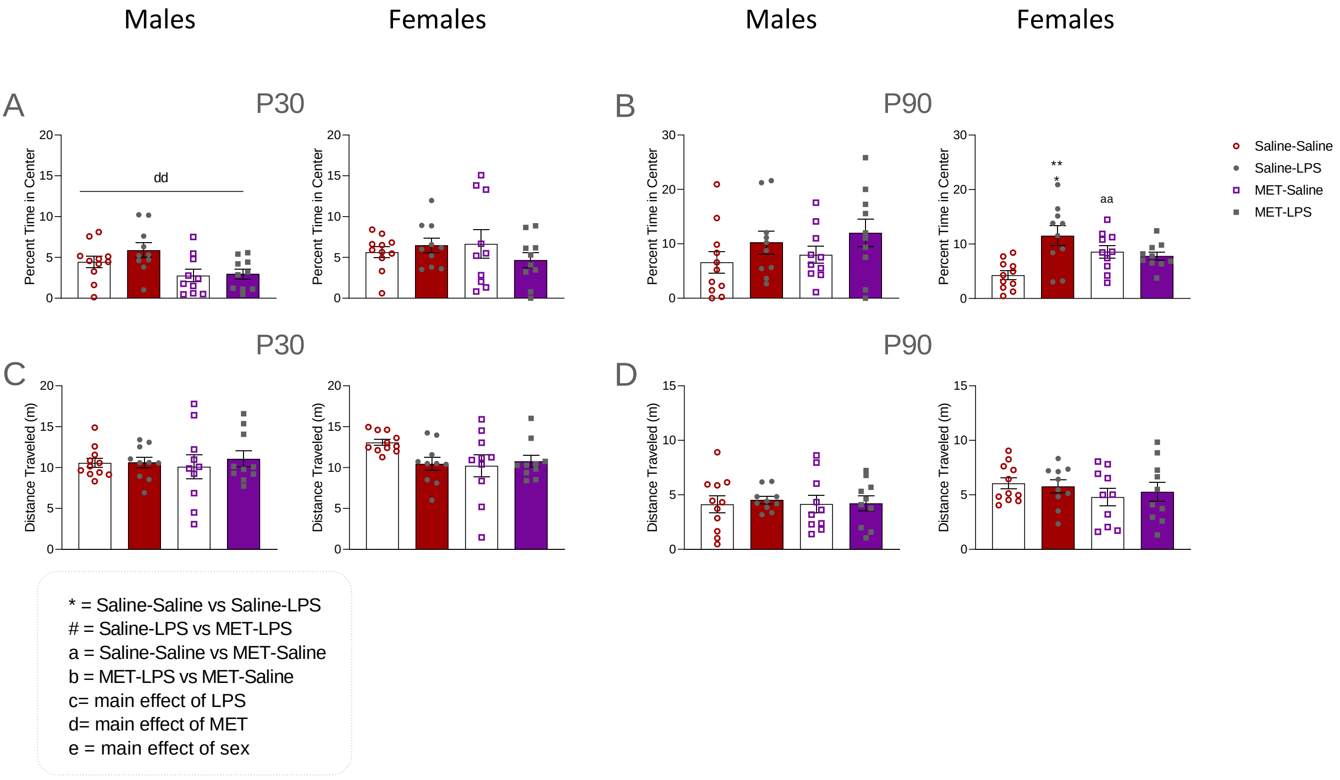

**Supplementary Results Figure 4. Maternal immune activation (MIA) and metyrapone (MET) induce sex-specific transcriptional effects in the ventral hippocampus. A** Threshold-free rank-rank hypergeometric overlap (RRHO) analysis revealed distinct transcriptional effects in males and females exposed to lipopolysaccharide (LPS). **B** RRHO revealed distinct transcriptional effects in males and females exposed to MET.

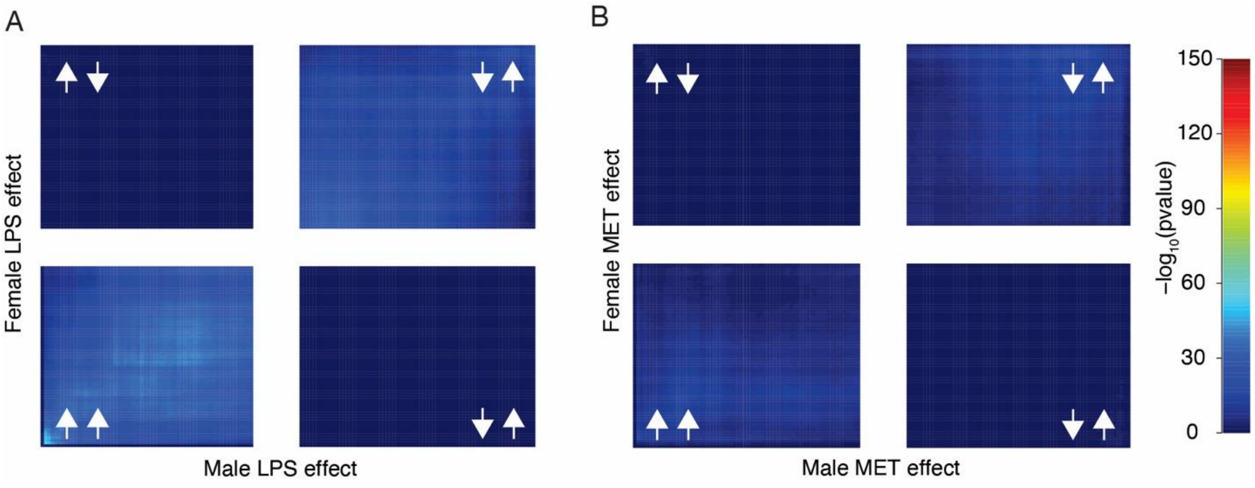


.

**Supplemental Results Figure 5. Effect of maternal immune activation (MIA) and metyrapone (MET) on transcriptomic analyses of ventral hippocampus tissue samples collected from adult male offspring.** Heatmaps of inflammation-, microglia-, LPS binding-, oxytocin-, epigenetics-, prolactin-, parvalbumin-, aldosterone/MR signaling-, glucocorticoid signaling-, GR binding-related genes, with effects represented as fold change (FC) between saline-saline and saline-LPS-exposed males, and between saline-LPS and MET-LPS-exposed males. Data are expressed as * FC > 1.3, ** p < 0.05, or *** p < 0.05 and FC > 1.3. LPS – lipopolysaccharide.

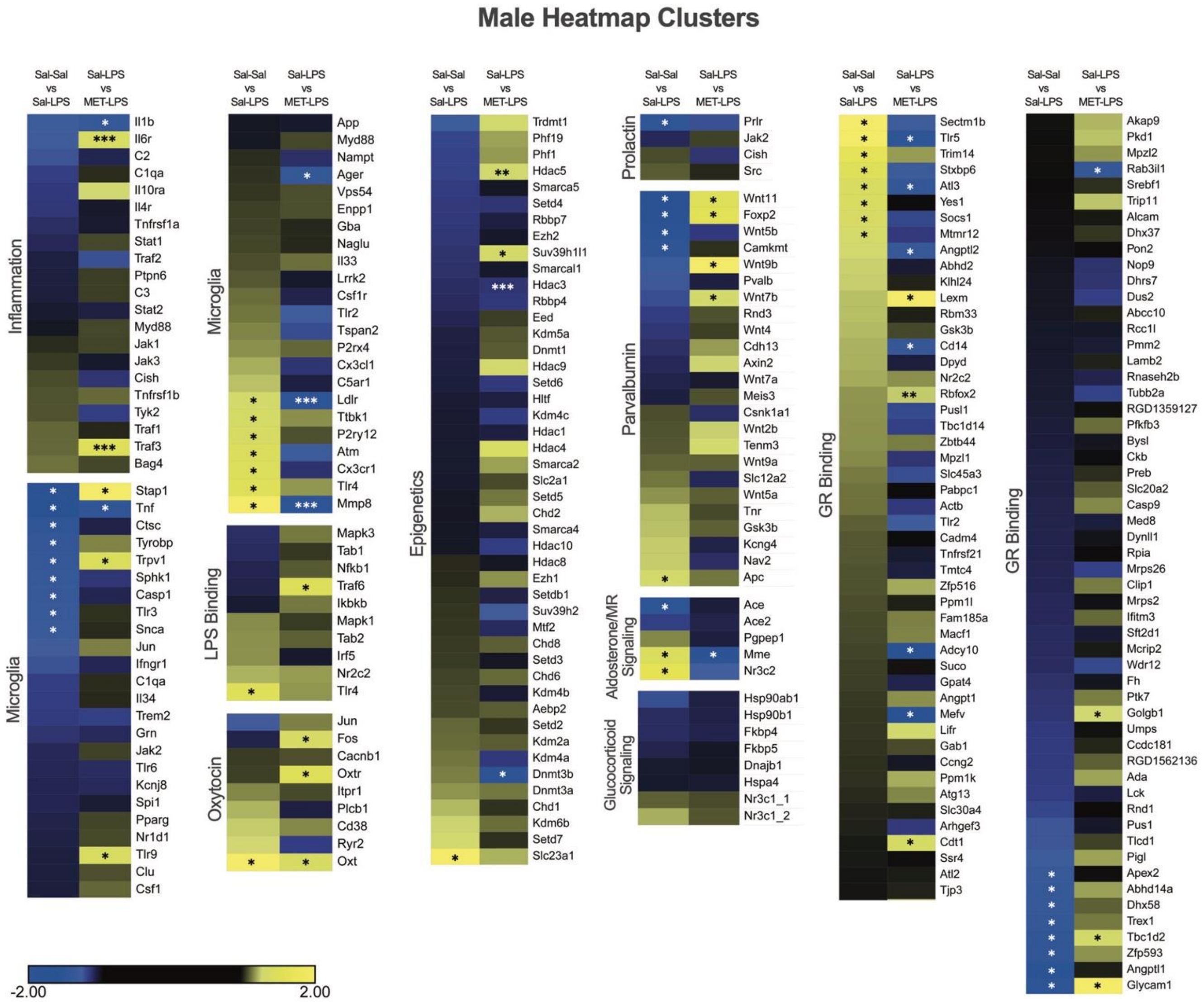

**Supplemental Results Figure 6. Effect of maternal immune activation (MIA) and metyrapone (MET) on transcriptomic analyses of ventral hippocampus tissue samples collected from adult female offspring.** Heatmaps of inflammation-, microglia-, LPS binding-, oxytocin-, epigenetics-, prolactin-, parvalbumin-, aldosterone/MR signaling-, glucocorticoid signaling-, GR binding-related genes, with effects represented as fold change (FC) between saline-saline and saline-LPS exposed females, and between saline-LPS and MET-LPS-exposed females. Data are expressed as * FC > 1.3, ** p < 0.05, or *** p < 0.05 and FC > 1.3. LPS – lipopolysaccharide.

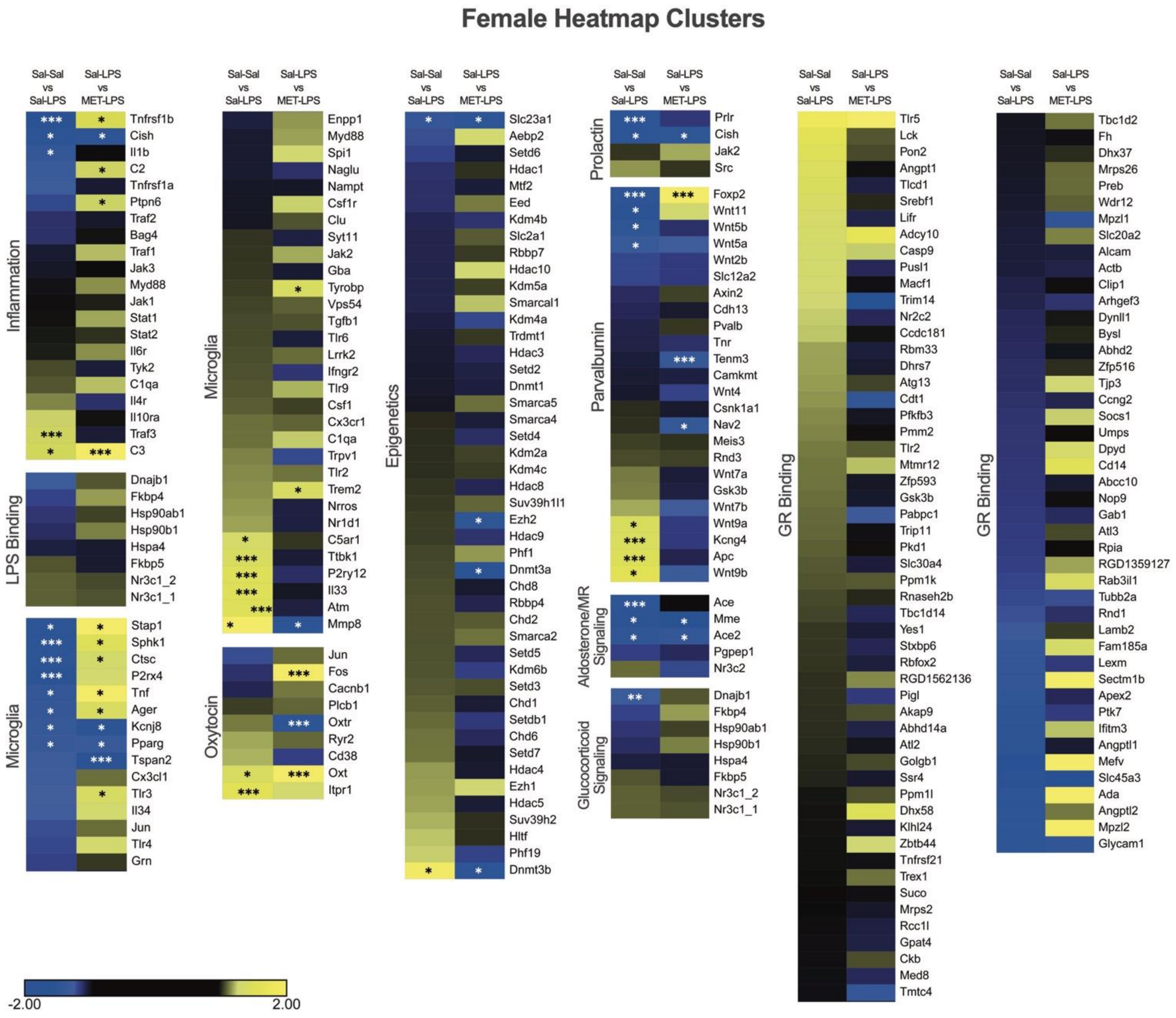

**Supplemental Results Table 3.** Summary of Gene Expression Patterns in Heatmap Clusters

| Heatmap Cluster | **MALE** Saline-Saline vs Saline-LPS | **MALE** Saline-LPS vs MET-LPS | **FEMALE** Saline-Saline vs Saline-LPS | **FEMALE** Saline-LPS vs MET-LPS |
|---|---|---|---|---|
| Inflammation | = | ↑ | ↑↓ | ↑ |
| Microglia | = | ↓ | ↑↓ | ↓ |
| LPS Binding | = | = | = | = |
| Epigenetics | = | ↑ | = | = |
| Parvalbumin | = | = | ↑↓ | ↑↓ |
| Glutamate/ GABA | ↑ | ↑↓ | ↑ | ↓ |
| Prolactin | = | = | ↓ | = |
| Oxytocin | = | = | ↑ | ↑↓ |
| Dopamine | ↑ | ↓ | ↑ | ↑↓ |
| Serotonin | ↑ | ↑↓ | ↑↓ | ↑↓ |
| Aldosterone/ MR Signaling | = | = | ↓ | = |
| GR Binding | = | ↑ | = | = |
| Glucocorticoid Signaling | = | = | ↓ | = |

**Supplemental Results Figure 7. Effect of maternal immune activation (MIA) and metyrapone (MET) on western blot analysis of receptor expression in the ventral hippocampus of adult offspring.** Ventral hippocampal **A** glucocorticoid receptor (GR) and **B** corticotropin-releasing factor receptor 1 (CRFR1) densitometric ratios for male and female offspring on P94. Neither GR/ß-Actin ratios nor CRFR1/ß-Actin ratios were significantly different between groups. Data are expressed as mean ± SEM. LPS – lipopolysaccharide.

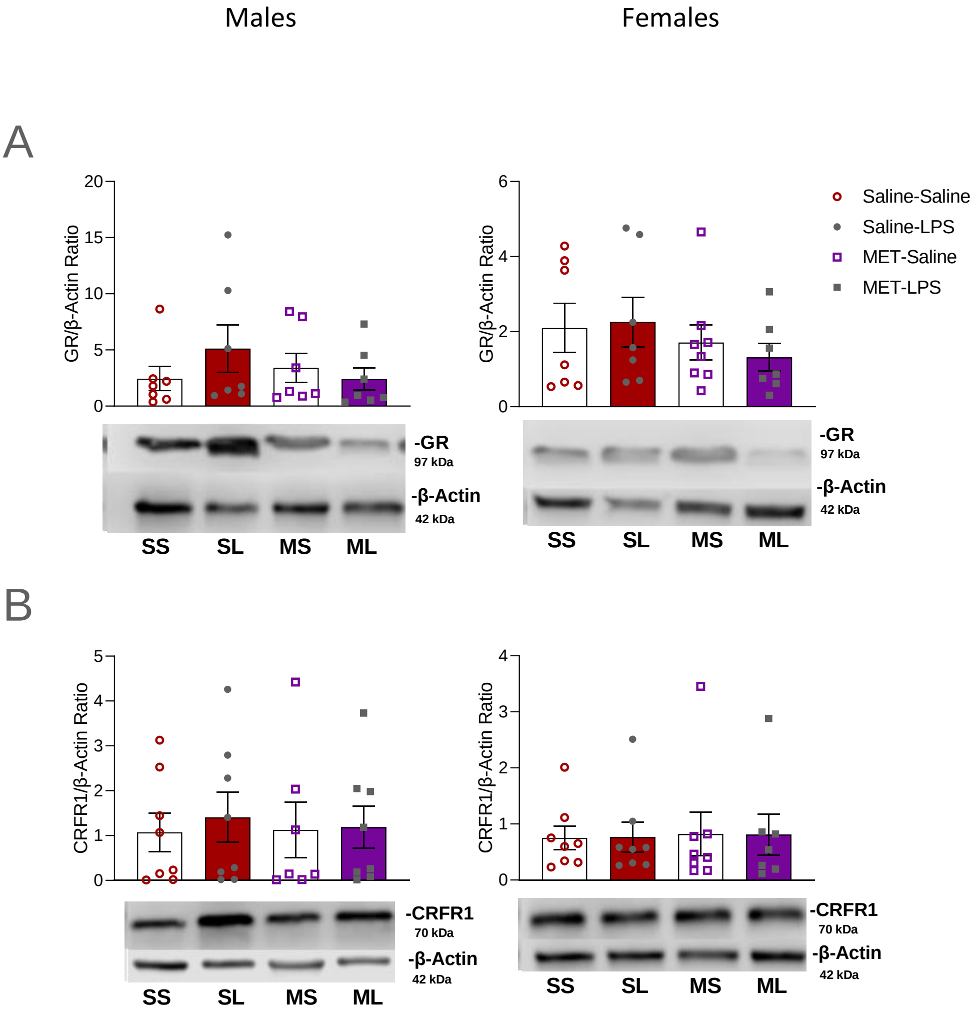
